# Nano-Mod-Amp reveals RNA sequence, structural and cell type specific features of pseudouridylation by PUS7

**DOI:** 10.1101/2025.10.30.685621

**Authors:** Rebecca Rodell, Ronit Jain, Hossein Shenasa, Matias Montes, Nicolas Robalin, Stefan Prodic, Eduardo Eyras, Nicole M. Martinez

## Abstract

Pseudouridines are abundant mRNA modifications that can impact splicing, translation, and stability to tune gene expression. PUS7 is one of the major mRNA pseudouridine synthase whose dysregulation leads to neurodevelopmental disorders and cancer, underscoring the critical function of PUS7-dependent pseudouridines. Beyond a short and degenerate consensus sequence, the molecular mechanisms underlying PUS7-mediated pseudouridylation remain unknown. A lack of targeted, high-throughput pseudouridine detection methods limits simultaneous interrogation of PUS7 regulatory features across many experimental conditions. We developed novel Nanopore sequencing tools, including Nano-Mod-Amp, to reveal pseudouridine stoichiometry, its RNA structural context, and dependence on PUS7 levels at specific sites across biological conditions. We identified a novel RNA structural signature that is associated with more efficient mRNA modification by PUS7. Pseudouridines are largely responsive to modulations in PUS7 protein levels, demonstrating the regulatory potential of varying PUS7 levels across cellular conditions. Conversely, PUS7 activity is also regulated in a cell-type specific manner, independent of PUS7 expression levels in a manner consistent with regulation by RNA structure and RNA binding proteins. Together, we developed Nanopore sequencing tools and uncovered new mechanisms of PUS7 regulation with a framework that can be applied to other RNA-modifying enzymes to query the regulation of the epitranscriptome.

**Highlights:**

- Nanopore direct RNA sequencing identifies PUS7-dependent pseudouridines with stoichiometry.
- Nano-Mod-Amp quantifies PUS7-dependent pseudouridines at hundreds of sites in high-throughput.
- MPRAs define RNA sequence and structural features associated with modification by PUS7.
- Individual PUS7 target pseudouridines are substoichiometric and poised for regulation.
- PUS7 activity is regulated by cell type in the absence of differences in PUS7 protein levels.

## Introduction

RNA modifications are chemical alterations to the canonical nucleosides that regulate many aspects of the RNA life cycle to impact fate and function^1^. Recent advances in transcriptome-wide RNA modification sequencing methods have determined the presence and locations of about ten distinct modifications in mRNAs^2–8^. Pseudourdine (Ψ), an isomer of uridine, is among the most abundant mRNA modifications^5–7^. Pseudouridines in mRNA can tune gene expression through impacts on splicing, translation, and mRNA decay^6,9–15^. Thirteen pseudouridine synthases (PUS) catalyze the isomerization of uridine to pseudouridine in humans. Over half of the pseudouridine synthases have been implicated in human disease, underscoring their importance to human health^1^.

PUS7 is one of the PUS with the most tRNA and mRNA targets ^5,6,10,12,16–18^. PUS7 modifies pre-mRNAs and has been shown to impact pre-mRNA processing, mRNA levels and translation ^6,10,19^. Predicted loss of function and missense mutations in the catalytic domain of PUS7 lead to neurodevelopmental disorders and have been linked to age-related macular degeneration^20–26^, while aberrant overexpression of PUS7 promotes glioblastoma tumorigenesis in a manner dependent on PUS7’s catalytic activity^17^. The underlying molecular mechanisms leading to PUS7 dysregulation and their link to disease outcomes remain to be determined. In addition to its significance in the context of disease, pseudouridylation of mRNAs by PUS7 is also altered in response to changing cellular conditions such as heat shock, metal ion stress, and lead exposure^6,19,27^,. These observations indicate that PUS7 activity is critical for proper gene expression during development, is dysregulated in disease, and may be regulated in diverse cellular contexts. However, what features, conditions, and mechanisms regulate human PUS7 activity are not well understood.

PUS7 mRNA targets are enriched for a UNUAR sequence motif, but only <5% of such motifs are pseudouridylated transcriptome-wide^10,28^, suggesting additional determinants of specificity. Although human PUS7 targets are enriched with the UNUAR motif, the relative importance of each nucleotide in the motif to the catalytic efficiency of PUS7 has not been systematically interrogated. Additional molecular features such as RNA structure and RNA-binding proteins are likely to influence PUS7 activity but have yet to be systematically investigated. While some studies of individual PUS7 targets suggest that RNA structure is important for modification of tRNA and non-coding RNAs^29,30^, another study suggests RNA structure is less critical for modification of an mRNA tested with yeast Pus7^31^. It is worth noting that PUS7 can modify non-coding RNAs in different 2D and 3D RNA structures, suggesting that PUS7 can modify within distinct and complex RNA structural contexts ^30^. Thus a unifying RNA structural motif and its impact on PUS7-dependent pseudouridine deposition remained to be elucidated.

Recently, multiple sequencing methods that rely on orthogonal modalities significantly advanced the transcriptome-wide study of pseudouridine and its stoichiometry (fraction of RNA molecules containing pseudouridines at a given position)^5–7,12,16,32^ ^33,34^. However, a challenge to studying the regulation and function of PUS7 is that these tools require high sequencing depth to achieve single-nucleotide quantification and robust statistical analysis of pseudouridines of interest in mRNAs. Moreover, high-sequencing costs limit the number of biological conditions that can be examined simultaneously. Existing targeted approaches for pseudouridine quantification are low throughput^35^, and lack the single molecule resolution afforded by long-read technologies such as Nanopore sequencing.

Here, we developed and applied orthogonal Nanopore sequencing tools for pseudouridine quantification, including Nanopore modification detection by amplicon sequencing (Nano-Mod-Amp), to uncover novel PUS7 regulatory features. We comprehensively characterize the mRNA targets of human PUS7, endogenously and with Massively Parallel Reporter Assays (MPRA) *in vitro* and *in cellulo*. We identified mRNA pseudouridines responsive to changes in PUS7 abundance, indicating sites are poised for regulation by enzyme concentration. However, PUS7 activity is not solely driven by sequence and enzyme levels but is influenced by additional layers of regulatory features which determine mRNA pseudouridylation. We found that PUS7 activity is associated with RNA structural features and influenced by the cellular environment. We established UGUAG as the version of the consensus UNUAR motif with the highest levels of PUS7-dependent pseudouridylation. We uncovered a surprising degree of structural heterogeneity among the experimentally probed 2D structures of human PUS7 mRNA targets yet discovered a unifying novel RNA structural signature among mRNA targets. We also demonstrate the impacts of cellular environment on regulation of PUS7-dependent pseudouridine by identifying a subset of pseudouridines that are cell-type specific that may be driven by RNA binding proteins and/or RNA structural remodeling. Our framework and approaches are broadly applicable to study the regulation and function of other RNA modifications and RNA modifying enzymes.

## Results

### Nanopore direct RNA sequencing reveals PUS7-dependent pseudouridine sites with stoichiometry

To characterize PUS7 regulatory features, we set out to identify PUS7 mRNA targets *de novo* using Nanopore direct RNA sequencing. In Nanopore sequencing, nucleotide identity is inferred from changes in current as RNA translocates through the pore with an applied voltage. Previous studies have demonstrated that pseudouridines lead to U-to-C mismatches during basecalling of Nanopore direct RNA sequencing^33,34^. Here, we assign individual pseudouridines to PUS7, through enzyme knockdown, and estimate absolute stoichiometry, based on synthetic standards containing all possible PUS7 recognition motifs (UNUAR; n=8; Figure 1A).^34,36^.

**Figure 1:**
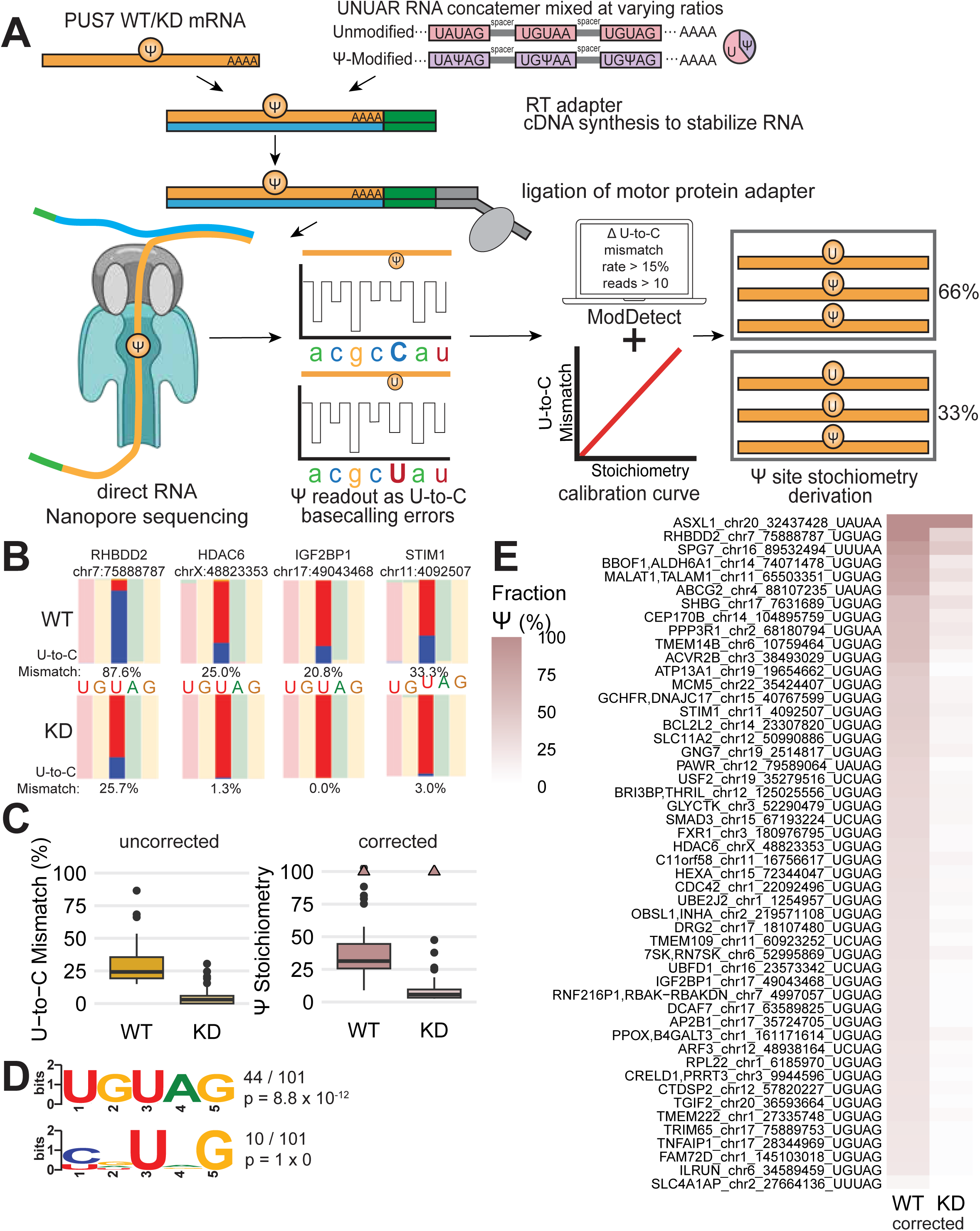
Nanopore direct RNA sequencing identifies PUS7-dependent pseudouridine with stoichiometry. (A) Schematic overview of Nanopore direct RNA sequencing. Total RNA from HepG2 WT, PUS7 KD and an unmodified and pseudouridine modified synthetic RNA standard with the 8 variants of the PUS7 motif is subjected to Nanopore’s direct RNA sequencing workflow. Ligation of an RT adapter is dependent on complementarity to a polyA tail to allow for cDNA synthesis to stabilize the RNA strand and unfold RNA structure. The RNA is threaded through the Nanopore by the motor protein and sequenced in the 3’ to 5’ orientation. (B) Genome browser style views of 4 representative PUS7-dependent sites from direct RNA sequencing observed in at least two out of three biological replicates with read coverage > 10. Nucleotide frequency is shown on the y-axis. Color-coded by nucleotide: blue, C; red, U; green, A; yellow, G. The pseudouridine is centered in the window. (C) (left) Uncorrected average U-to-C mismatch rate for 50 PUS7-dependent sites identified by direct RNA sequencing in at least two out of three biological replicates with read coverage > 10 and mismatch rate > 15% in a UNUAR motif. (right) Calculated pseudouridine stoichiometry for 50 PUS7-dependent sites identified in left panel based on calibration curves from UNUAR standard. Triangles indicate corrected stoichiometry greater than 100% for one site. (D) Top three motifs from STREME analysis of 101 identified PUS7-dependent sites from direct RNA sequencing with coverage > 10 and differential U-to-C mismatch rate > 15% between PUS7 KD and WT in two out of three replicates compared to background set of uridines with coverage > 10. Abbreviations: WT, wild-type; KD, dox-inducible knockdown (E) Corrected pseudouridine stoichiometry in WT and KD of PUS7-dependent sites identified from direct RNA sequencing in at least two out of three replicates, with coverage > 10 and differential U-to-C mismatch rate > 15% between PUS7 KD and WT in UNUAR motifs. Corrected values determined from average U-to-C mismatch rate and UNUAR standard calibration curves. Sites are labeled with genomic location and motif containing the pseudouridine.

To benchmark the detection of pseudouridine, we sequenced all 8 variants of the known PUS7 sequence motif in an unmodified synthetic RNA concatemer (UNUAR) and a site specifically modified RNA (UNΨAR) (Supplemental Figure 1A). We found that pseudouridine in each PUS7 motif presents as U-to-C mismatches, demonstrating the suitability of this approach for identifying PUS7 targets. (Supplemental Figure 1B,C). To derive absolute stoichiometries, we mixed the modified and unmodified standard at defined ratios and generated calibration curves (Supplemental Figure 1D). Some motifs (UGUAG) display high correlation between U-to-C mismatches and pseudouridine stoichiometry, while others (UAUAA) are underestimated due to sequence context dependent effects. This variance in U-to-C mismatch rate by sequence motif highlights the need for synthetic standards to derive absolute stoichiometry.

To identify PUS7-dependent mRNA pseudouridines, we sequenced total RNA from wild-type (WT) and PUS7 knockdown (KD) HepG2 cells. The standard Nanopore direct RNA sequencing protocol selectively captures mRNA^10,34^ To assign PUS7-dependent pseudouridines, we required a U-to-C mismatch difference ≥ 15% between PUS7 WT and KD samples and a per site read number ≥ 10, with reproducibility in two out of three biological replicates (Figure 1A). Because our replicates ranged in coverage (∼2-6M total reads per sample in line with the expectation of direct RNA flow cells), insufficient coverage in a subset of replicates is a limiting factor for detection of PUS7-dependent pseudouridines. Thus, we also included sites observed in one replicate and also reported in other transcriptome-wide datasets^10,12,16^. Our pipeline identified 101 candidate PUS7 pseudouridine mRNA sites (Figure 1B,C) with an enrichment^37^ of the PUS7 UGUAG motif in alignment with previous studies^37^ (Figure 1D, Table S1). The 50 mRNA pseudouridines within the UNUAR that are dependent on PUS7 include *RHBDD2*, *HDAC6*, *IGF2BP1* and *STIM1* (Figure 1B, Table S1). PUS7-sensitive positions outside the UNUAR motif could represent pseudouridines in an imperfect consensus context or RNA modifications indirectly impacted by PUS7 knockdown.

Previous work has shown that pseudouridines occur across mRNA without strong enrichment in a particular transcript region (e.g. 5’UTR, CDS, 3’UTR)^9^. We asked whether identified PUS7-dependent pseudouridines in mRNAs were enriched in specific transcript regions. The distribution of PUS7 mRNA targets across transcript features is skewed towards 3’ UTRs, similar to all detected uridines, suggesting that the skewed distribution is a byproduct of the 3’ coverage bias due to Nanopore sequencing in the 3’ to 5’ direction. Our analysis suggests that PUS7 modifies across transcript regions (Supplemental Figure 1F).

We used our UNUAR calibration curves to derive absolute stoichiometry of PUS7 mRNA targets in the consensus UNUAR motif. Corrected pseudouridine stoichiometry for PUS7 mRNA targets range from 8% to 100% with a median of 31%, demonstrating that many PUS7 mRNA targets are highly modified (Figure 1C,E, Supplemental Figure 1E). Our synthetic standard and calibration curves now enable deriving quantitative pseudouridine stoichiometries from Nanopore direct RNA sequencing data. These advances allow comparing stoichiometry across sequence context and relating modification levels to downstream biological functions.^9^ In summary, our Nanopore direct RNA sequencing pipeline identifies PUS7 mRNA targets with absolute stoichiometry. The key additions of a genetic control and synthetic standards to the workflow enable quantification of pseudouridine stoichiometries at sites assigned to a specific PUS in a generalizable approach.

### Nano-BID-amp orthogonally validates endogenous modification and PUS7-dependency

Nanopore direct RNA sequencing is a powerful approach for *de novo* discovery of RNA modifications transcriptome-wide. However, epitranscriptomic research requires orthogonal validation of candidate RNA modification sites due to coverage limitations and method specific biases. We developed Nano-Mod-Amp as a versatile high-throughput approach to validate candidate sites and study their regulation across diverse conditions.

Many RNA modifications can be read out indirectly, following chemical labeling, as mutations introduced during reverse transcription. Nano-Mod-Amp is a modular targeted Nanopore approach to read out mutations that result from various chemical treatments of RNA through cDNA sequencing. Nano-Mod-Amp enables cost effective, high coverage detection of endogenous pseudouridine with long-reads in medium to high-throughput. Compared to transcriptome-wide approaches such as Nanopore direct RNA sequencing, where coverage per mRNA ranges from tens to hundreds of reads, our targeted approach selectively amplifies target mRNAs of interest with thousands of reads per mRNA. The per-sample barcodes allow for the pooling of amplicons from many biological conditions. This method allows going from RNA to analyzed Nanopore sequencing data in as little as 2 days. For reverse-transcription, we use template-specific primers with a 10 nucleotide Unique Molecular Identifier (UMI) and an adapter for compatibility with Nanopore cDNA ligation barcoding sequencing (Figure 2A). Sequential bead-based clean-up removes UMI-containing RT primer to enable accurate PCR duplicate collapsing. A gene-specific forward primer adds the second adapter for compatibility with Nanopore’s PCR barcoding primers. Nanopore motor protein adapters are then ligated for sequencing.

**Figure 2:**
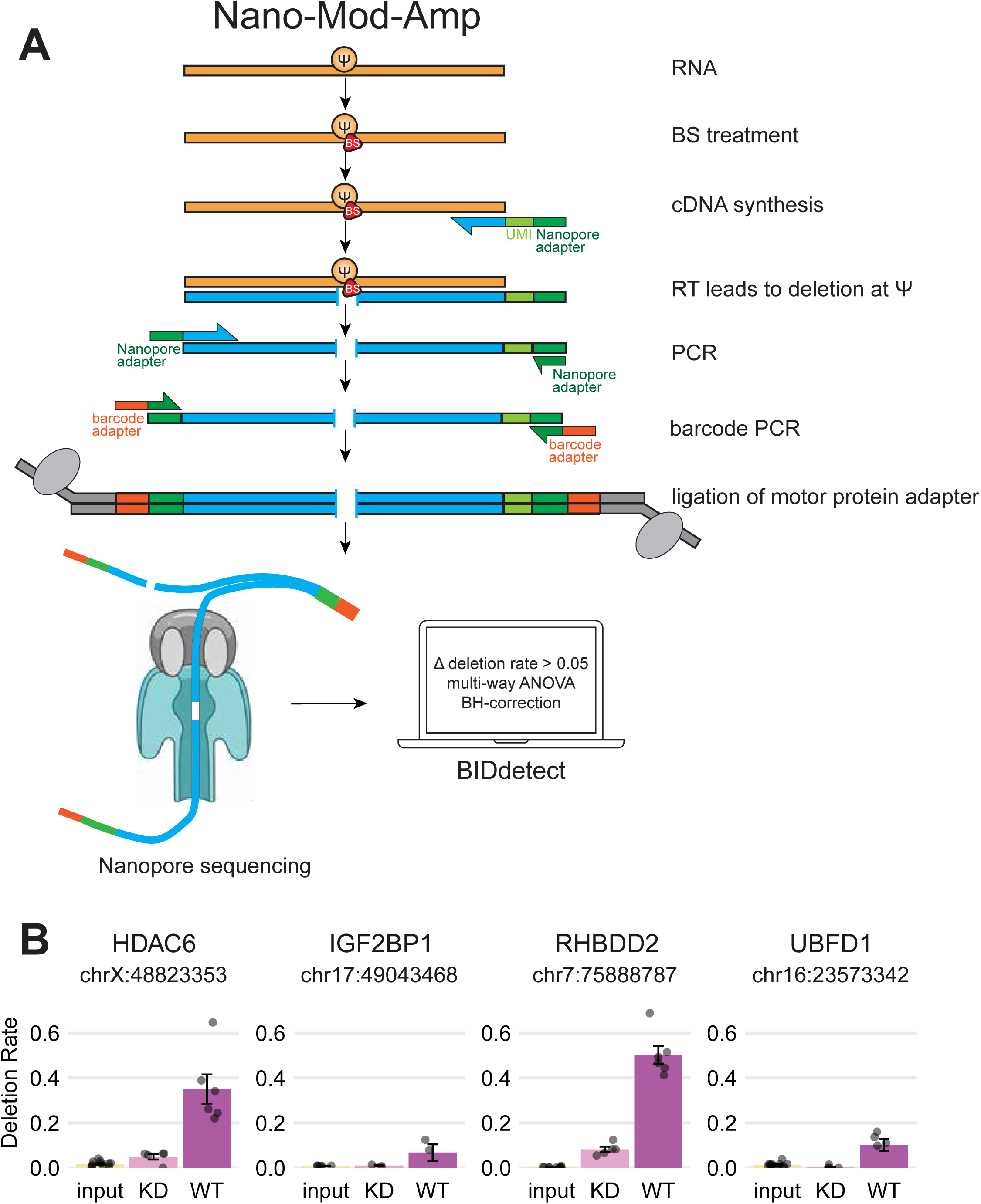
Nano-BID-amp orthogonally validates endogenous modification and PUS7-dependency. (A) Schematic overview of Nano-Mod-Amp sequencing approach with bisulfite labeling of pseudouridines (Nano-BID-Amp). RNA is treated with BS, labeling the pseudouridine. During cDNA synthesis with amplicon-specific primers including a UMI and Nanopore adapter, the pseudouridine-BS adduct creates a deletion. PCR amplification with amplicon-specific primers including Nanopore adapters is followed by Nanopore barcode PCR, ligation of motor protein adapter, and cDNA Nanopore sequencing. BIDdetect analyzes data to identify sites with >20 reads of coverage, > 0.05 Δ deletion rate between treatment and control groups, and p < 0.05 from multi-way ANOVA with Benjamini-Hochberg correction applied. (B) Endogenous Nano-BID-Amp from HepG2 and HEK293T cells validate PUS7-dependent pseudouridine with a Δ deletion rate > 0.05 between PUS7 WT and KD with a Benjamini-Hochberg-corrected p-value less than 0.05 (n = 6, multi-way ANOVA). Abbreviations: BS, bisulfite, WT, wild-type; KD, dox-inducible knockdown

We applied Nano-Mod-Amp to orthogonally assess candidate PUS7-dependent pseudouridine sites we identified through Nanopore direct RNA sequencing. Bisulfite treatment of RNA followed by desulphonation leads to deletions at pseudouridines during reverse transcription^12,16^ that we read out with Nano-BID-Amp^12,16^. In contrast to Nanopore direct RNA sequencing, BID treatment leads to more uniform deletion rates across sequence contexts and does not require a synthetic standard calibration curve to compare across sequence contexts^12^. Importantly, we obtained comparable deletion rates at pseudouridines in the context of a synthetic RNA standard by sequencing amplicons using the described Nanopore library preparation strategy or Illumina sequencing (Supplemental Figure 2A). The targeted nature of our long-read amplicon sequencing approach produces high read-depth at low cost, enabling powerful tests of statistical significance. We developed a complementary computational pipeline to quantify mutational signatures resulting from native RNA modifications or from chemical treatments. RNA structure or sequence context can impact mutational signatures. Our statistical framework considers how these factors impact chemical treatment in the context of a biological perturbation of interest (cell type, enzyme level, etc.) across replicates.

We assessed PUS7-dependence among pseudouridines identified through Nanopore direct RNA sequencing, by applying Nano-BID-Amp to polyA-selected mRNA from WT or PUS7 KD HepG2 (Figure 2B). We validated PUS7 dependence for 5 randomly selected endogenous amplicons (300 – 880 nts long), including *HDAC6*, *IGF2BP1*, *RHBDD2*, and *UBFD1* (Figure 2B, Table S3), supporting the accuracy of our approach. We observe BID-dependent deletions corresponding to pseudouridine levels that decrease significantly following PUS7 KD (Figure 2B). Altogether, Nano-BID-Amp is a simple and robust method for targeted yet high-throughput quantification of pseudouridines across conditions. Nano-BID-Amp validates pseudouridines identified by Nanopore direct RNA sequencing or other transcriptome-wide approaches.

### Massively parallel reporter assays recapitulate PUS7-dependent pseudouridines

While modification by PUS7 has been studied under basal conditions, regulation in response to changes in enzyme concentration or trans-acting factors has not been investigated. To interrogate multiple regulatory features simultaneously in a single reaction, we designed a massively parallel reporter assay (MPRA)^10,28,38^. We used Nano-BID-Amp to measure PUS7 activity *in vitro* and *in cellulo* across hundreds of targets and in many experimental conditions (Figure 3A). We included endogenous PUS7 mRNA targets identified by direct RNA sequencing (Figure 1) and expanded this list to include additional annotated PUS7 mRNA sites (521 total), as well as 254 PUS1 mRNA targets identified in other studies as negative controls ^10,12,16^. We extracted 130 nucleotides of flanking sequence centered on the pseudouridine and appended 5’ and 3’ adapters as handles for Nano-BID-Amp. We included a T7 promoter upstream of the 5’ adapter to transcribe the MPRA pool *in vitro.* For *in cellulo* experiments, we cloned the MPRA pool into the 3’ UTR of a mCherry reporter under a Pol II promoter (Figure 3A).

**Figure 3:**
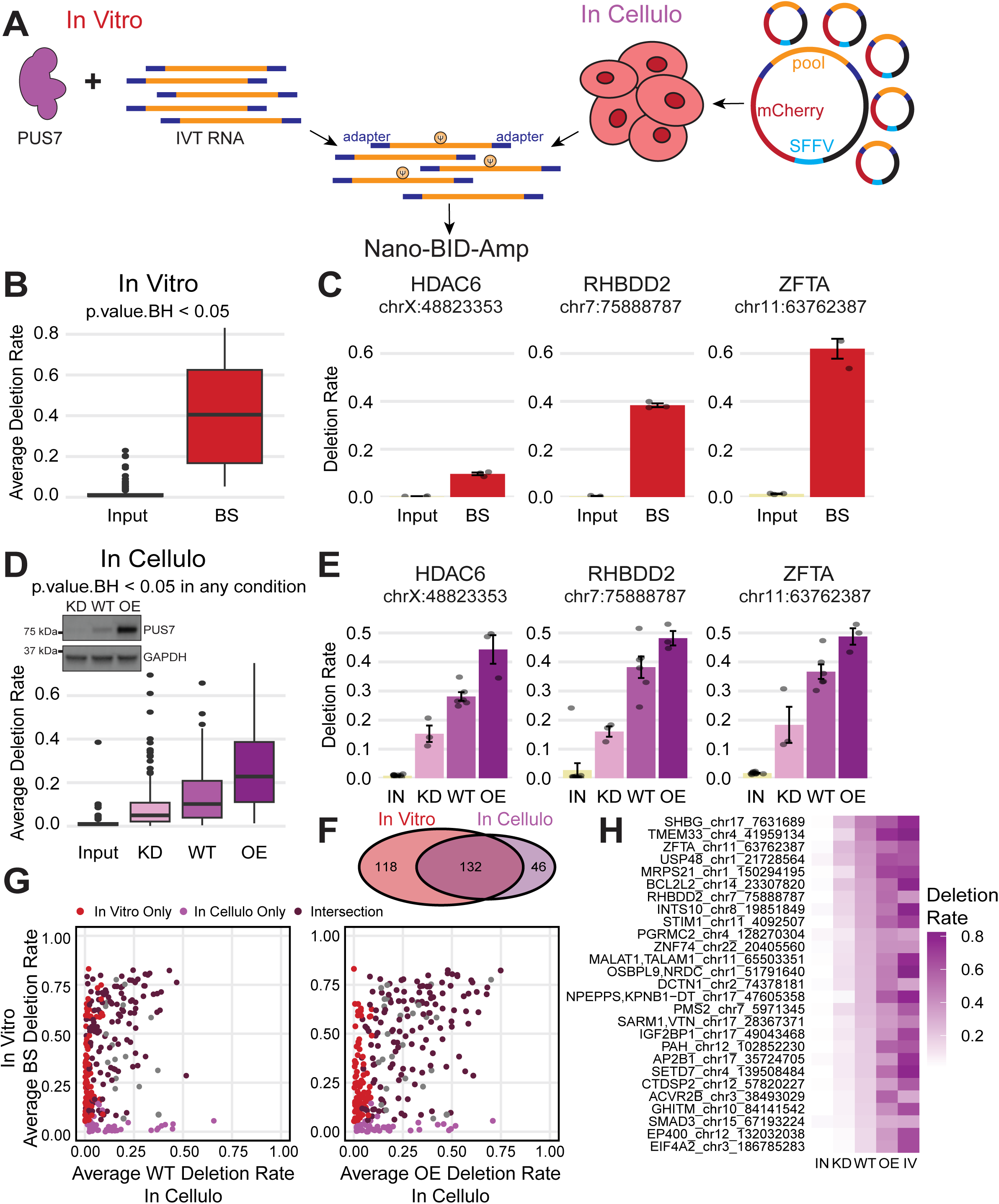
Massively parallel reporter assays reveal differences between *in vitro* and *in cellulo* PUS7-sensitivity. (A) Schematic of *in vitro* and *in cellulo* MPRA. For *in vitro* Nano-BID-Amp MPRA, recombinant PUS7 was incubated with *in vitro* transcribed and folded RNA to allow *in vitro* pseudouridylation. For *in cellulo* Nano-BID-Amp MPRA, a plasmid containing the pool in the 3’ UTR of mCherry downstream of a SFFV Pol II promoter was transiently transfected into cells to allow pseudouridylation. Cells were collected 48 hours later, RNA was extracted. Both sets of RNA were then used in the Nano-BID-Amp workflow with universal handles from the 5’ and 3’ adapters to quantify extent pseudouridylation based on bisulfite induced deletions. (B) *In vitro* pseudouridylation PUS7-targets in MPRA with recombinant PUS7 identified 250 sites through Nano-BID-Amp with a Δ deletion rate > 0.05 between BS and input and Benjamini-Hochberg-corrected p-value less than 0.05 (n = 3, multi-way ANOVA). (C) Representative individual PUS7-dependent pseudouridine from *in vitro* pseudouridylation with Nano-BID-Amp. Sites had a Δ deletion rate > 0.05 between BS and input and Benjamini-Hochberg-corrected p-value < 0.05 (n = 3, multi-way ANOVA). (D) Representative Western blot of PUS7 levels in WT, KD and OE HepG2 cells with GAPDH as a loading control. Transient transfection of PUS7-target MPRA into HepG2 and HEK293T cells with KD, WT, and OE PUS7 levels identified 178 sites through Nano-BID-Amp with a Δ deletion rate > 0.05 and Benjamini-Hochberg-corrected p-valued less than 0.05 (n > 3, multi-way ANOVA). These 178 sites have a statistically significant different deletion rate between WT and KD and / or WT and OE in at least one cell type. (E) Representative individual PUS7-dependent pseudouridine from *in cellulo* pseudouridylation with Nano-BID-Amp in HEK293T cells. Sites had a Δ deletion rate > 0.05 and Benjamini-Hochberg-corrected p-value < 0.05 (n = 3, multi-way ANOVA). (F) Overlap of statistically significant sites from *in vitro* and *in cellulo* Nano-BID-Amp MPRA. Sites had a Δ deletion rate > 0.05 and Benjamini-Hochberg-corrected p-valued < 0.05 (n = 3, multi-way ANOVA). (G) Comparison of average deletion rates between *in vitro* MPRA and *in cellulo* MPRA WT (left) and PUS7 OE (right) for PUS7-dependent sites. Light purple – significant *in cellulo* only; red – significant *in vitro* only; dark purple – significant *in vitro* and *in cellulo*; grey – significant in one condition with uncertainty in the other condition masking statistical significance. Significance based on a Benjamini-Hochberg-adjusted p-value less than 0.05 (n = 3, multi-way ANOVA). (H) Deletion rates of significant sites modified *in vitro* and *in cellulo* in Nano-BID-Amp MPRA with a Δ deletion rate > 0.1 from KD to WT and WT to OE. Deletion rates for *in cellulo* and *in vitro* experiments are shown. Abbreviations: KD, constitutive knock down; WT, wild-type; OE, overexpression; BS, bisulfite treatment; IV, in vitro; IN, input

To measure modification *in vitro*, we performed *in vitro* transcription and pseudouridylation with recombinant PUS7^10,39,40^ (Supplemental Figure 3A), followed by Nano-BID-Amp in three biological replicates. We detected 250 pseudouridines with a > 0.05 difference in deletion rate between input and bisulfite-treated samples that were statistically significant (Figure 3B, Table S4). This included recapitulation of 80% of PUS7-dependent pseudouridine in UNUAR motifs that we identified by Nanopore direct RNA sequencing (Figure 1B-D). *HDAC6*, *RHBDD2*, and *ZFTA* have representative pseudouridines recapitulated *in vitro* with recombinant PUS7 (Figure 3C).

To measure pseudouridylation in the context of the *in cellulo* MPRA, we transiently expressed the cloned MPRA into two cell lines: HepG2 and HEK293T. Pseudouridine levels in the heterologous *in cellulo* MPRA context are comparable to levels measured at endogenous loci (Figure 2B, 3E). The MPRAs demonstrate that 130 nucleotides of sequence surrounding the pseudouridine are sufficient to recapitulate endogenous modification by PUS7 both *in vitro* and *in cellulo*. To determine how PUS7 concentration impacts pseudouridylation, we transfected the MPRA in WT, PUS7 KD, and PUS7-V5 overexpression (OE) cells and performed Nano-BID-Amp (Figure 3D). To establish PUS7-dependence, we looked for a statistically significant difference in pseudouridine levels between WT and KD or OE when HEK293T and HepG2 data were analyzed together or separately. We detected 178 PUS7-dependent pseudouridines in at least one condition (Figure 3D, Supplemental Figure 3B, Table S5). The *in cellulo* MPRA recapitulated 74% of PUS7-dependent pseudouridine that we identified by Nanopore direct RNA sequencing (Figure 1B-D). The majority of pseudouridines are modulated by PUS7 KD and PUS7 OE (Figure 3D,E). In contrast, some sites reached apparent modification saturation with WT PUS7 levels as modification did not increase with PUS7 OE compared to WT levels (Supplemental Figure 3C). Other sites that were undetectable in WT were modified upon PUS7 overexpression (Supplemental Figure 3D). These results demonstrate that tuning PUS7 protein levels can be a strategy for regulated pseudouridylation across biological conditions. Collectively, we recapitulate PUS7 mRNA targets identified by Nanopore direct RNA sequencing with *in vitro* and *in cellulo* MPRAs and demonstrate the modulation of PUS7-dependent pseudouridine in response to changes in PUS7 concentration. We also show that MPRAs can be used to study how sequence, structure and other cellular factors influence PUS7 pseudouridine levels with Nano-BID-Amp.

### Massively parallel reporter assays reveal differences between in vitro and in cellulo PUS7-sensitivity

While in the *in vitro* MPRAs, pseudouridylation is dictated by sequence and enzyme levels alone, the *in cellulo* environment introduces trans-acting factors and spatiotemporal impacts on pseduouridylation. To understand how the cellular context shapes pseudouridine levels, we compared pseudouridines *in vitro* versus *in cellulo*. We found 132 pseudouridines that are PUS7-dependent both *in vitro* and *in cellulo*, 118 pseudouridines were significantly modified to detectable levels only *in vitro*, and 46 sites were significantly modified only *in cellulo* (Figure 3F). These results suggest that factors within the cellular environment such as co-transcriptional recruitment of PUS7, compartmentalization, RNA binding proteins (RBPs), and/or dynamic RNA structures play a significant role in setting the PUS7-dependent pseudouridine landscape in cells. In contrast, the lack of these variables *in vitro* and an excess of PUS7 could allow for greater modification, both in terms of total number of sites and modification stoichiometry (Figure 3G,H). Thus, the cellular environment is essential for a subset of PUS7 sites which cannot be efficiently pseudouriydylated even with vast excess of enzyme *in vitro*.

### PUS7 mRNA targets share sequence and structural features despite heterogeneous 2D structures

Sequence motifs drive the binding^41–43^ and catalysis of many RBPs and RNA modifying enzymes^44^. PUS7-dependent pseudouridines occur in the context of a UNΨAR sequence motif^10,28^, which we recapitulate in our transcriptomic direct RNA data, *in vitro* MPRA and *in cellulo* MPRA (Figure 1, Supplemental Figure 4A). However, less than 1% of UNUAR motifs detected in our Nanopore direct RNA sequencing data (Figure 1) are pseudouridylated. In our MPRAs, we see almost no PUS7-dependent sites modified outside of this motif, but not every UNUAR motif is modified (Table S3). Even *in vitro*, where only RNA and recombinant PUS7 are present, UNUAR is not sufficient for robust modification in all cases. RNA structure is the only other feature present *in vitro* that could influence modification. Distinct RNA structural motifs are necessary for pseudouridylation of mRNAs by yeast Pus1^28^ and human TRUB1^38^, which led us to hypothesize that RNA structure influences PUS7-mediated pseudouridylation. We leveraged our comprehensive, cross-validated list of PUS7 mRNA targets that are sensitive to PUS7 protein levels (Figure 1-3) to uncover RNA structural features associated with modification.

To identify shared RNA structural features among PUS7 mRNA targets, we performed *in vitro* RNA structure probing with the selective 2’-hydorxyl acylation analyzed by primer extension (SHAPE) reagent 2-methylnicotiniv acid imidazolide (NAI)^45^ using our Nano-Mod-Amp workflow. SHAPE reagents react with the 2’OH of flexible or unpaired RNA nucleotides. In SHAPE-MaP, reverse transcriptase misincorporates at SHAPE adducts during cDNA synthesis in the presence of manganese-containing buffers^46^. The resulting mutations are read out by sequencing and subsequently converted to SHAPE reactivities. Normalized SHAPE reactivities can be used for experimentally informed RNA structure predictions. We took advantage of the modular workflow of Nano-Mod-Amp (Figure 4A) and replaced BID with SHAPE treatment, (Nano-SHAPE-Amp) and otherwise subjected the MPRA to the same library prep protocol. Nano-SHAPE-Amp facilitates long-read, high-throughput, targeted RNA structure probing.

**Figure 4:**
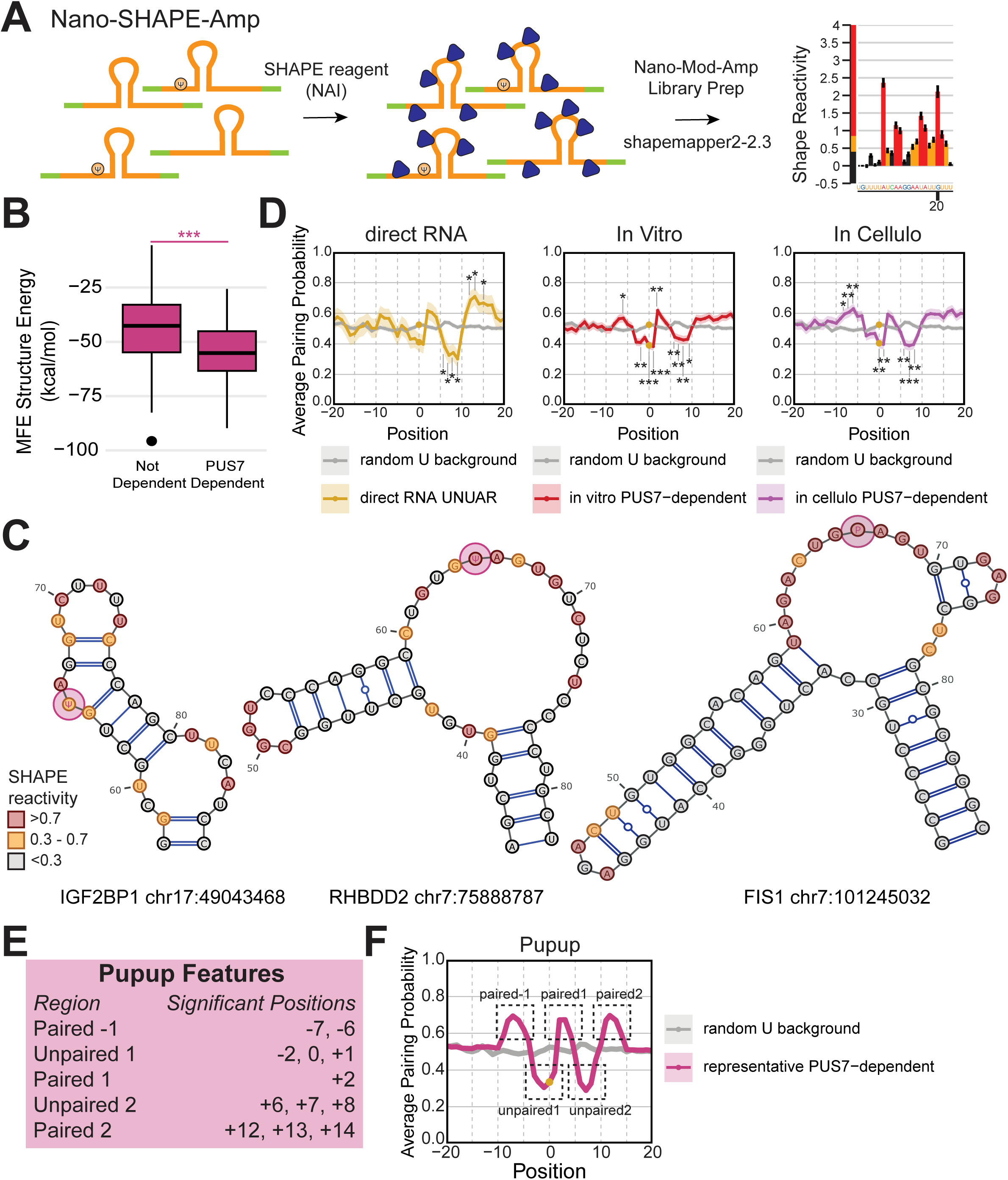
PUS7 mRNA targets share sequence and structural features despite heterogeneous 2D structures. (A) Schematic of Nano-SHAPE-Amp, wherein RNA is treated with SHAPE reagent NAI to form adducts are unpaired nucleotides. Nano-Mod-Amp workflow is followed. During cDNA synthesis, SHAPE adducts cause mutational signatures. Sequencing data is deduplicated and fed into shapemapper2 to determine SHAPE reactivity profiles. (B) Minimum free energy was calculated for MPRA sequences using SHAPE-constrained RNAfold. MPRA sequences not dependent on PUS7 *in vitro* or *in cellulo* with Nano-BID-Amp were compared to sequences found to be PUS7-dependent *in vitro* and *in cellulo* with Nano-BID-Amp. Mann-Whitney U test with Benjamini-Hochberg corrected p-value = 6.87 x 10^-14^. (C) Representative MEA structures determined with SHAPE-constrained RNAfold are shown of PUS7-dependent pseudouridine identified *in vitro* and *in cellulo* Nano-BID-Amp MPRA (132 in total). (D) Metaplots of pairing probabilities calculated by SHAPE-constrained RNAfold for PUS7-dependent pseudouridine identified in (left) direct RNA, (middle) MPRA *in vitro*, and (right) MPRA *in cellulo*. Background pairing probabilities centered around a random U (MPRA) are represented in grey, PUS7-target pairing probabilities are represented by the colored lines. Position 0 (gold dot) represents the pseudouridine. Sites where the pseudouridine was not determinable to a single position were excluded from these and subsequent analysis. Welch’s T-test with Benjamini-Hochberg correction was applied to all sites in the +/-15 nt window to determine statistically significant difference. (E) Features of the Pupup signature, defined by statistical significance between background and PUS7-targets at the specified sites in panel C. (F) Generalized schematic of how the regions that make up the Pupup signature in pairing probability metaplots. Notes: * p < 0.05; ** p < 0.01, *** p < 0.001

Nano-SHAPE-Amp reactivity data from our MPRA enables experimentally-informed predictions of the 2D RNA structures present among PUS7 mRNA targets^47^ (Figure 4A). SHAPE-informed folding of PUS7 mRNA targets using RNAfold mirrored purely *in silico* predictions (Figure 4B-D, Supplemental Figure 4B-C). To determine if PUS7 targets are more structured, we compared the free energy of folding between PUS7 targets and non-targets and found that PUS7 targets have a significantly lower free energy (Figure 4B). Our systematic approach revealed that the SHAPE-informed 2D maximal expected accuracy (MEA) structures adopted by PUS7 mRNA targets are heterogenous (Figure 4C).

To determine whether there are any unifying features of the diverse structures adopted by PUS7 mRNA targets, we made metaplots of the average pairing probabilities and tested for statistically significant differences relative to a randomly sampled background uridine distribution (Figure 4D). Across the direct RNA and MPRA PUS7-dependent sites, a paired-unpaired-paired-unpaired-paired signature arose, with the pseudouridine falling in the first unpaired region, which we termed the “Pupup” signature (Figure 4E,F). This signature is unique to PUS7 targets as it is not present among unmodified UNUAR sequences (Supplemental Figure 4C). *In silico* folding only predicted the downstream features of the Pupup signature, emphasizing the importance of experimentally probing RNA structure as opposed to relying on *in silico* modeling alone (Supplemental Figure 4C). Comparing individual PUS7 mRNA target MEA structures to the Pupup signature revealed full (*IGF2BP1*) and partial matches. *RHBDD2* conforms to the first Pup, where the pseudouridine is in an unpaired loop flanked by the first two paired segments of Pupup. *FIS1* conforms to the downstream upup, where an unpaired pseudouridine is followed by a downstream paired-unpaired-paired segment (Figure 4C). The Pupup signature provides a unifying model for the recognition of heterogeneous RNA structures of PUS7 mRNA targets, though not all targets conform to all features of the signature.

Our analysis of PUS7 mRNA targets confirmed the enrichment of the UNUAR sequence motif and identified a novel RNA structure signature among otherwise heterogenous 2D structures. We applied the Nano-Mod-Amp workflow to SHAPE-based chemical probing and determined experimentally informed RNA structures of PUS7 mRNA targets *in vitro*. The presence of the Pupup motif potentially explains why PUS7 does not modify all UNUAR sites, as RNA structure is associated with pseudouridylation.

### The UGUAG motif and Pupup signature are associated with higher PUS7-dependent pseudouridine stoichiometry

To further dissect the features associated with PUS7 modification, we asked how sequence and structure varied with pseudouridylation level. To do this, we binned *in vitro* and *in cellulo* PUS7-dependent sites into four quartiles, where Q1 represents the least modified sites and Q4 represents the most highly modified sites (Figure 5A). We observed a strong preference for the UGUAG sequence motif over UNUAR in the most highly modified (Q4) compared to the most lowly modified (Q1) sites, implying UGUAG is most amenable to modification (Figure 5B, Supplemental Figure 5A). Lowly modified Q1 sites showed a UNUAG motif, indicating flexibility is most tolerated at the –1 position albeit at the expense of lower modification levels. We find a lower free energy of folding among the PUS7 mRNA target structures with higher levels of modification, suggesting that more highly modified sites have a higher average thermodynamic stability (Supplemental Figure 5B). We observed differences in the pairing probabilities across quartiles, with Q4 sites more strongly conforming to the Pupup signature (Figure 5C, Supplemental Figure 5C). Strikingly, we also noticed a statistically significant decrease in the pairing probability of the pseudouridine residue between Q1 and Q4 (Figure 5D), suggesting higher possibility of modification when the target U is more accessible to PUS7. In summary, higher PUS7-dependent pseudouridylation is associated with an unpaired pseudouridine in a UGUAG sequence context and an RNA structural context that conforms to the Pupup signature.

**Figure 5:**
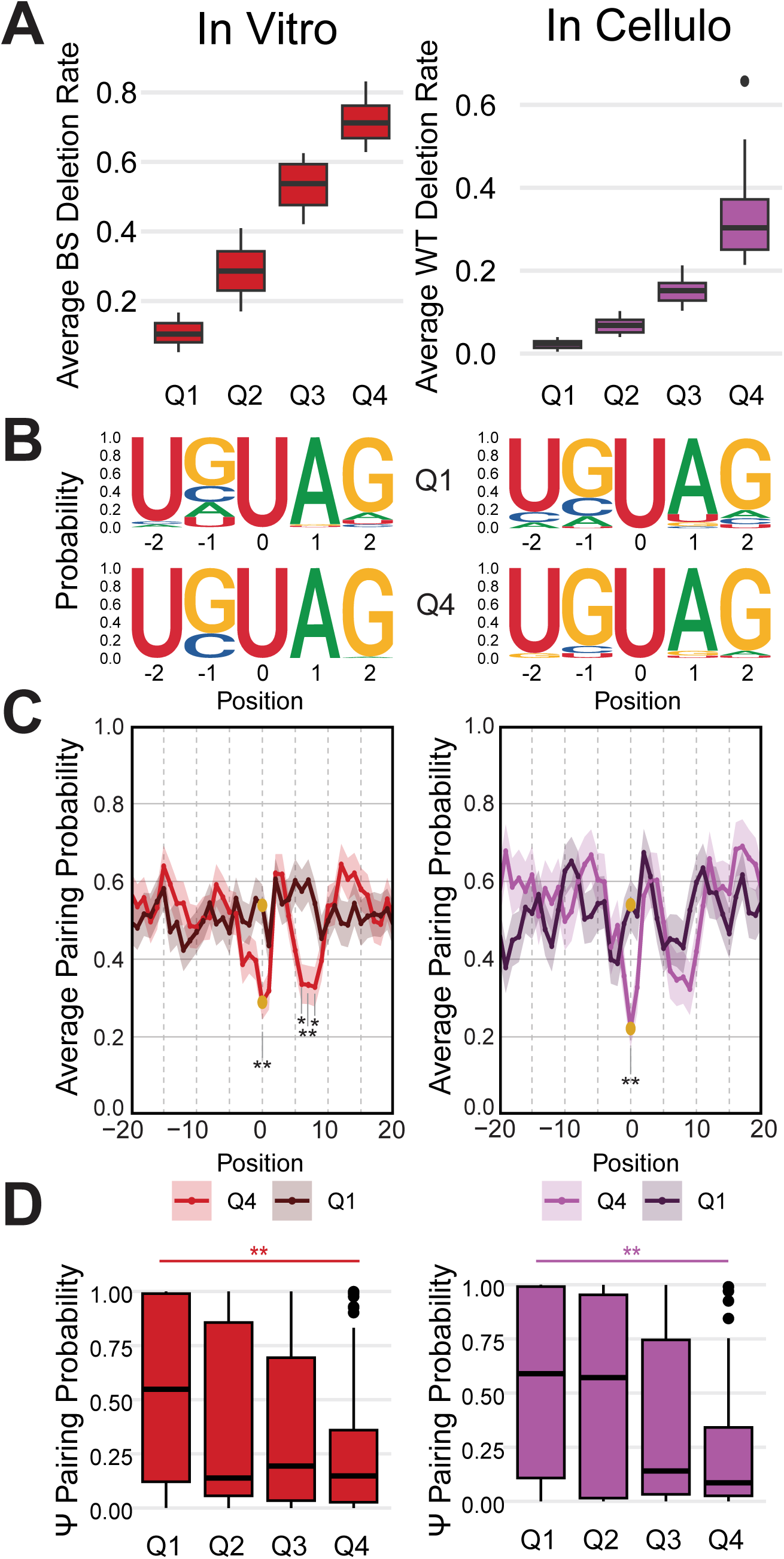
The UGUAG and Pupup motif are associated with higher PUS7-dependent pseudouridine stoichiometry. (A) Average deletion rate of (left) *in vitro* and (right) *in cellulo* PUS7-dependent pseudouridine identified *in vitro* and *in cellulo* with Nano-BID-Amp MPRA divided into quartiles. Q1 represents the least modified sites and Q4 the most modified sites. BS-treated deletion rates are shown for *in vitro*; WT BS-treated deletion rates are shown for *in cellulo*. (B) Metaplots of nucleotide frequencies for PUS7-dependent pseudouridine from Q1 (top) and Q4 (bottom) identified in (left) MPRA *in vitro* Nano-BID-Amp and (right) MPRA *in cellulo* Nano-BID-Amp. Position 0 represents the pseudouridine. Sites where the pseudouridine was not determinable to a single position were excluded from these and subsequent analysis. (C) Metaplots of pairing probabilities calculated with SHAPE-constrained RNAfold for PUS7-dependent pseudouridine from Q1 and Q4 identified in (left) MPRA *in vitro and* (right) MPRA *in cellulo*. Position 0 (gold dot) represents the pseudouridine. Welch’s T-test with Benjamini-Hochberg correction was applied to all sites in the +/-15 nt window to determine statistically significant difference. (D) Pairing probabilities calculated with SHAPE-constrained RNAfold for the pseudouridine position for quartiles, as defined in A, for PUS7-dependent sites identified in (left) MPRA *in vitro* and (right) MPRA *in cellulo* Nano-BID-Amp. Wilcoxon rank-sum test p-value of 0.046 (*in vitro*) and 0.006 (*in cellulo*) for difference between Q1 and Q4 pairing probability. Abbreviations: Q1, first quartile; Q2, second quartile; Q3, third quartile; Q4, fourth quartile; BS, bisulfite-treated; WT, wild-type Notes: * p < 0.05; ** p < 0.01, *** p < 0.001

### PUS7-dependent pseudouridines are differentially regulated by cell type

Pseudouridine profiles are altered in response to stressors^6,19,34^ and vary across mouse tissues^12^, suggesting condition specific regulation. The *in cellulo* MPRA enabled us to interrogate PUS7-dependent pseudouridine for differential modification between human cell lines of different tissue origins: HepG2 (liver) and HEK293T (kidney). Many PUS7-dependent pseudouridines showed no cell type specificity (Figure 6A). Our analysis revealed 49 PUS7 target sites with higher modification in HEK293T than HepG2 (Figure 6 A-C, Supplemental Figure 6A,B). We validated the differential PUS7-dependent pseudouridylation by cell type using endogenous Nano-BID-Amp. While both *RHBDD2* and *HDAC6* are PUS7-dependent (Figure 2B), *HDAC6* is more highly modified in HEK293T compared to HepG2 cells while *RHBDD2* is not (Figure 6D). These findings demonstrate cell type specific regulation of pseudouridines by PUS7.

**Figure 6:**
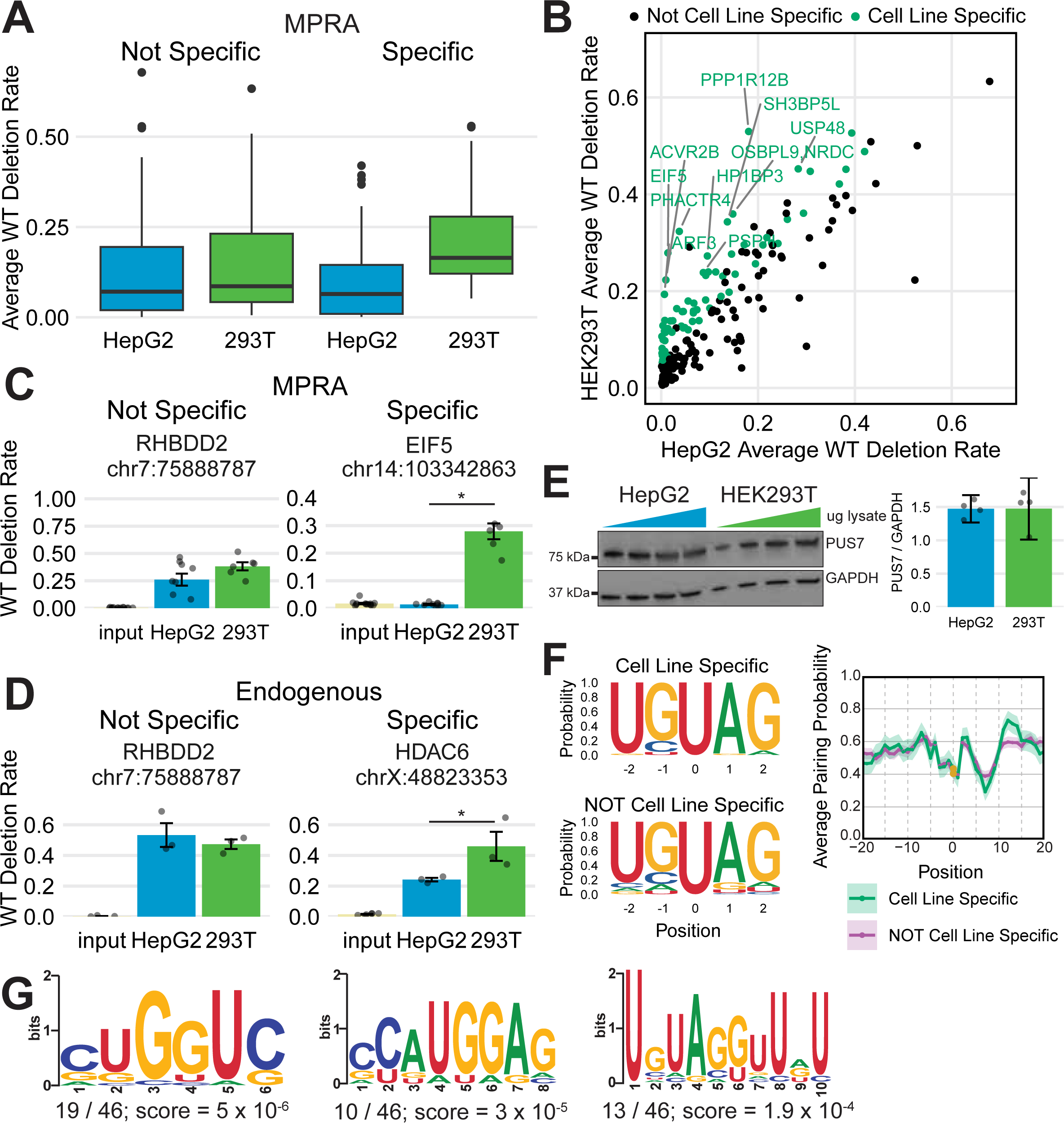
PUS7-dependent pseudouridines are differentially regulated by cell type. (A) Transient transfection of PUS7-target MPRA into HepG2 and HEK293T WT cells revealed 49 sites with a Δ deletion rate > 0.05 between HepG2 and HEK293T in Nano-BID-Amp and a Benjamini-Hochberg-adjusted p-value < 0.05 (n = 3, multi-way ANOVA). (B) Deletion rates in WT *in cellulo* MPRA Nano-BID-Amp for 178 PUS7-dependent sites compared by cell type. The 49 PUS7-dependent cell type specific pseudouridine with a Δ deletion rate > 0.05 between HepG2 and HEK293T and a Benjamini-Hochberg-adjusted p-value less than 0.05 (n = 3, multi-way ANOVA) are highlighted. (C) Representative cell type specific PUS7-dependent sites from *in cellulo* MPRA Nano-BID-Amp with a Δ deletion rate > 0.05 between HepG2 and HEK293T cells and Benjamini-Hochberg-corrected p-value less than 0.05 (n = 3, multi-way ANOVA). (D) Endogenous Nano-BID-Amp from HEK293T and HepG2 validate cell type specific PUS7-dependent pseudouridine with a Δ deletion rate > 0.05 between HepG2 and HEK293T cells and Benjamini-Hochberg-corrected p-valued less than 0.05 (n = 3, multi-way ANOVA). (E) Western blot of varying amount of HepG2 and HEK293T RIPA lysates (left) and quantification, normalizing PUS7 levels to GAPDH expression (right) with anti-PUS7 Abcam antibody. (F) (left) Nucleotide frequency metaplots for PUS7-dependent cell-line specific (top) and PUS7-dependent, not cell type specific (bottom) between HepG2 and HEK293T as determined by *in cellulo* MPRA Nano-BID-Amp. (right) Metaplots of pairing probabilities calculated by SHAPE-informed RNAfold for PUS7-dependent cell type specific and not cell type specific sites. Position 0 (gold) represents the pseudouridine. Sites where pseudouridine was not determinable to a single position were excluded from these analyses. Welch’s T-test with Benjamini-Hochberg correction was applied to all sites in the +/-15 nt window to determine statistically significant difference, and none were found Top three STREME consensus motifs for PUS7-dependent cell type specific sites enriched compared to non-PUS7-dependent cell-line specific sites, as identified in *in cellulo* MPRA Nano-BID-Amp. Abbreviations: WT, wild-type; Massively Parallel Reporter Assay (MPRA).

To investigate the mechanisms behind the cell-type specificity of this subset of PUS7-dependent pseudouridines, we followed multiple lines of inquiry. RBP levels, RBP localization, RNA decay and export rates and RNA structure, among other factors, vary with cell type^48–52^ and could tune pseudouridine levels. First, we demonstrated that PUS7 proteins levels are equivalent between cell types (Figure 6E). PUS7 localization, visualized through immunofluorescence, also does not substantially differ between the cell types (Supplemental Figure 6C). Next, we compared sequence and structure motifs between cell-type specific and non-specific PUS7-dependent pseudouridines. Both sets of pseudouridines similarly conformed to the UNUAG sequence motif and Pupup structural signature (Figure 6F). We then asked whether trans-acting RBPs might recruit PUS7 or compete with PUS7 to access mRNA target sites. Using STREME^37^, we uncovered a CUGGUC motif, which is associated with the CELF and MBNL family of RBPs, enriched in 19 PUS7-dependent cell-line specific sequences over PUS7-dependent sequences lacking cell-line specificity (Figure 6G). These results suggest that RBP-mediated recruitment of or competition with PUS7 could be a potential mechanism for cell-type specificity.

## Discussion

Prior work identified PUS7 as a major pseudouridine synthase acting on mRNA, yet the full repertoire of PUS7 mRNA targets remains to be determined. Furthermore, why certain mRNAs sites are modified by PUS7 while others are not remains an open question. Obtaining answers to these questions has been limited by the lack of quantitative tools to measure pseudouridine fraction in a site-specific manner across many experimental conditions. Here, we used complementary pseudouridine quantification methods, direct RNA sequencing and Nano-Mod-Amp, to comprehensively identify novel features associated with basal and condition-specific mRNA pseudouridylation by PUS7.

### Direct RNA sequencing identifies PUS7 mRNA targets and pseudouridine stoichiometry

We implemented a Nanopore direct RNA sequencing pipeline to identify PUS7-dependent mRNA pseudouridines transcriptome-wide.^33,34,53^. We demonstrate that genetic assignment to PUS7 based on differential U-to-C mismatches between enzyme depleted and control samples can be used to robustly identify PUS7 mRNA targets with high validation rates using Nano-Mod-Amp as an orthogonal approach. We generated a synthetic RNA standard with all eight PUS7 UNUAR sequence motifs to benchmark U-to-C mismatch rates, making it possible to derive pseudouridine stoichiometries from Nanopore direct RNA sequencing data at PUS7-dependent mRNA pseudouridines. Our Nanopore direct RNA sequencing framework is a powerful approach to identify and determine the stoichiometry of PUS7 mRNA targets *de novo* with quantitative resolution. Our Nanopore direct RNA sequencing data and standards will be a valuable resource for training PUS7-centric pseudouridine basecallers that can estimate pseudouridine stoichiometry from raw signal in the future.

### Nano-Mod-Amp as a new tool for studies of the epistructurome

Nanopore direct RNA sequencing allows for *de novo* discovery of PUS7-dependent pseudouridines transcriptome-wide. However, these experiments are expensive and limited by coverage, making it difficult to quantify changes in pseudouridine levels across many experimental conditions. By developing Nano-Mod-Amp, we addressed these limitations and established an orthogonal validation approach for pseudouridines at known or predicted locations. Nano-Mod-Amp now enables targeted, cost effective, quick-turnaround, high coverage, and statistically principled quantification of pseudouridines multiplexed across many biological conditions. Nanopore sequencing libraries can be performed in house or sent out to a commercial vendor. In Nano-BID-Amp, pseudouridines are derivatized with bisulfite^12,16^ and are quantified as deletions during cDNA synthesis. We illustrate the modularity of the Nano-Mod-Amp approach by swapping out the bisulfite for other chemical probes such as those that report on RNA structure (e.g. SHAPE reagents). In principle, the Nano-Mod-Amp approach can be applied to any modification, natural or artificial, that produces mutational signatures during cDNA synthesis. For example, Nano-Mod-Amp could be combined with solvent accessibility probes that report on RBP binding^54^ to quantify protein-agnostic changes in RBP binding across conditions. Our complementary computational pipeline quantifies diverse mutational profiles to report on statistically significant RNA features of interest. These new approaches will enable the field to elucidate the mechanisms of the condition-specific regulation for various modified nucleotides and their relationship to RNA structures.

### Sequence and new structural features associated with modification by human PUS7

Prior work has characterized sequence and structural features that determine pseudouridylation by human TRUB1 and yeast Pus1, yet a comprehensive understanding of the molecular basis for site selection by human PUS7 was lacking. Various studies^9^ have found a UNUAR motif enriched among PUS7 mRNA targets; however we find that a UNUAR motif alone is insufficient to confer specificity in cells or *in vitro* (Table S4 & S5). Studies on individual targets suggest that tRNA structural features are generally important for tRNA modification by PUS7^2930^. However, these studies are counter balanced by another which suggests that RNA structure may not be as important for modification of an mRNA target of yeast Pus7^31^. Thus, the importance of RNA secondary structure in PUS7 mediated pseudouridylation remained an open question.

Here, we systematically addressed this question for hundreds of human PUS7 mRNA targets with engineered MPRAs that are compatible with Nano-Mod-Amp. Our systems-level approach discovered several structural features that are associated with pseudouridylation of PUS7 mRNA targets: 1) stable structures with a lower free energy of folding, 2) an unpaired pseudouridine position that is more accessible, 3) enrichment of the Pupup structural signature compared to mRNAs that are not targets. The Pupup structural signature constitutes a unifying structural feature among the otherwise heterogeneous 2D structures adopted by PUS7 mRNA targets. In contrast, yeast Pus1 and human TRUB1 target mRNAs share homogeneous consensus structures consisting of a bulged stem loop where the pseudouridine position is at the base of the stem and a short stem loop where the pseudouridine position is in the loop respectively. These differences in RNA structural preferences among the studied pseudouridine synthases underscore the importance of characterizing the features that guide modification by each of the thirteen human PUS enzymes. This will inform assigning unique and redundant functions to paralogs of the same families. Our results suggest that RNA structure is important for binding and/or modification by PUS7 and that accessibility of the target uridine in a single stranded region facilitates pseudouridylation. The novel features that we define here will help better predict PUS7 mRNA targets from sequence, however we show that other factors within the cellular environment, such as trans-acting factors also likely influence pseudouridine levels. Our characterization of PUS7 targets will begin to lay the groundwork for dissecting the regulation and role of individual targets in developmental and disease contexts where PUS7 is critical^9^.

### Condition-specific regulation of PUS7 activity in cells

Whether PUS7 levels are saturating or changes in the cellular concentration of PUS7 are sufficient to drive changes in pseudouridylation had not been addressed. The increased expression of PUS7 could have no impact on targets if the levels of PUS7 are already saturating, lead to increases in pseudouridylation stoichiometry at sites that are modified basally, or lead to the pseuoduridylation of new or suboptimal sites. By varying the levels of PUS7, we find that many pseudouridines are titratable with changes in PUS7 protein level, and thus amenable to regulation by changes in PUS7 concentration that are observed naturally across cell types (Supplementary Figure 6D) and in disease contexts like in cancer compared to normal tissue^17^ (Supplemental Figure 6E). However, we also see examples where pseudouridine levels do not increase with higher levels of PUS7 and others that are detectably modified only when PUS7 is overexpressed. This suggests that pseudouridine sites that are not modified basally in a given cell type could be induced in the context of disease related overexpression.

Our high-throughput investigation of PUS7 mRNA pseudouridylation in two cell types revealed that regulation of PUS7 activity is also possible in the absence of differences in PUS7 protein levels or changes in localization. Interestingly, a subset of PUS7 mRNA targets are pseudouridylated to a higher degree in HEK293T cells compared to HepG2 cells. These results suggest that other features such as trans-acting RBPs and/or context specific RNA structure could underlie cell type specific regulatory mechanisms. Analysis of the identified cell-type specific sequences revealed a similar strength of the UNUAR sequence motif and similar intrinsic folding propensity between cell type specific and constitutive PUS7 mRNA targets (Figure 6F). We hypothesize that differential pseudouridylation by PUS7 between cell types could be explained by a cell type-specific RBP that competes with or recruits PUS7. Analysis of sequence motifs enriched among transcripts with cell type specific pseudouridines identified a CUG-motif primarily upstream of modified sites in support of these models. This motif is bound by members of the CELF and MBNL family of RBPs that function in pre-mRNA splicing and post-transcriptional gene regulation ^41,43,55–57^. Regulation of pseudouridines by RBPs could be reminiscent of splicing regulation through competition between RBPs binding to the same motif and recruitment of an RBP by another RBP bound to an adjacent motif.^58–60^ RNA structural remodeling has been observed in different cell states^61^ and cell types^48^ and could mechanistically explain cell type specific pseudouridines. Future work investigating whether RBPs and/or structural remodeling between cell types impact PUS7 activity will provide additional insight into the new mechanisms of regulation identified here. Together, our results reveal that regulation of PUS7 activity can be achieved in both a concentration dependent and independent manner, and that RNA structure and trans-acting factors may set the cell type specific pseudouridine landscape.

In conclusion, orthogonal Nanopore sequencing methods allowed us to identify PUS7-dependent pseudouridines and provide the most comprehensive and detailed characterization of human PUS7 targets and regulatory features to date. PUS7 mRNA targets fold into heterogeneous secondary structures but are characterized by adherence to a Pupup structural signature that is associated with higher modification. Our newly developed Nano-Mod-Amp is a versatile tool to quantify pseudouridines and other RNA modifications that can be detected by RT mutational signatures in high-throughput across diverse biological conditions. We used these approaches to uncover cell-line specific PUS7-dependent pseudouridines and provide some evidence consistent with regulation by differences in trans-acting factors and/or RNA structural remodeling in cells. Nano-Mod-Amp is broadly applicable across diverse RNA modifications to study the regulation of the epitranscriptome and its relationship with RNA secondary structure. Our work represents an important step in determining the mechanisms that could underly diseases resulting from dysregulation of PUS7 by characterizing mRNA targets and features that guide PUS7 activity. These PUS7 mRNA targets could represent new therapeutic targets, while our identified regulatory features pave the way for new modalities to control PUS7-dependent pseudouridylation in future work.

## Figure Legends

**Supplemental Figure 1:** Nanopore direct RNA sequencing benchmarking and calibration standards. Related to Figure 1. (A) Schematic of UNUAR concatemer in unmodified and fully-pseudouridylated states. Each of the 8 UNUAR motifs are separated by a 14-nucleotide spacer. A polyA tail is appended to the end of the sequence to allow for integration in Nanopore direct RNA sequencing pipelines. (B) Genome browser style views for UNUAR concatemer at each of the UNUAR motifs. Unmodified template on top, fully pseudouridylated template on bottom. Nucleotide frequency is shown on the y-axis. Color-coded by nucleotide: blue, C; red, U. The pseudouridine is centered in the window. (C) U-to-C mismatch rates across the UNUAR concatemer for the 8 U-Ψ mixes sequenced. Positions follow the order of the motif in A. Non-U positions are shown as having a mismatch rate of 0. (D) Calibration curves for each of the UNUAR motifs. UNØAR concentration is shown on the x-axis, observed U-to-C mismatch rate in direct RNA sequencing is shown on the y-axis. Line of best fit was determined through a linear model and fitted to each motif individually. (E) Uncorrected U-to-C mismatch rate and corrected Ψ stoichiometry in WT and KD of PUS7-dependent sites identified from direct RNA sequencing in at least two out of three replicates, with coverage > 10 and mismatch rate > 15% in a UNUAR motif. Corrected values determined from average U-to-C mismatch rate and UNUAR standard calibration curves. Sites are labeled with genomic location and motif containing the pseudouridine. (F) Distribution of PUS7-dependent pseudouridine identified in direct RNA sequencing with > 15% U-to-C mismatch rate across transcriptome regions and RNA type in comparison to background uridine distribution. All sites were observed in two out of three biological replicates and had > 10 reads of coverage.

**Supplemental Figure 2:** Comparison of deletion rates obtained by Nanopore compared to Illumina amplicon sequencing. Related to Figure 2. (A) Genome browser views of deletion rates for a fully pseudouridylated standard (with pseudouridine at 4 positions) from Nano-BID-Amp protocol with either Nanopore (top) or Illumina (bottom) sequencing.

**Supplemental Figure 3:** Investigation of PUS7-dependence with Massively parallel reporter assays *in vitro* and *in cellulo* related to Figure 3. (A) Coomassie stain of recombinant PUS7 purification. Fractions from all lanes shown under FPLC fractions were pooled and used as recombinant PUS7 protein for *in vitro* pseudouridylation. (B) Overlap of Nano-BID-Amp MPRA *in cellulo* PUS7-dependent sites. PUS7-dependent pseudouridine were identified from HEK293T data analyzed alone, HepG2 and HEK293T data analyzed together, and HepG2 data analyzed alone. PUS7-dependency was calculated as Δ deletion rate > 0.05 between WT and KD or WT and OE with a Benjamini-Hochberg-corrected p-value < 0.05 (n = 3, multi-way ANOVA). The union of all sites was taken to identify 178 PUS7-dependent sites used in further analyses. (C) Representative sites from Nano-BID-Amp MPRA in HEK293T cells with a significant difference between WT and PUS7 KD *in cellulo* (Δ deletion rate > 0.05, Benjamini-Hochberg-corrected p-value < 0.05 (n = 3, multi-way ANOVA)) and no significant different between WT and OE, indicating saturation at PUS7 WT levels. (D) Representative sites from Nano-BID-Amp MPRA in HEK293T cells with a significant difference between WT and PUS7 OE *in cellulo* (Δ deletion rate > 0.05, Benjamini-Hochberg-corrected p-value < 0.05 (n = 3, multi-way ANOVA)) and no significant different between WT and KD, indicating absence of modification at PUS7 WT levels.

**Supplemental Figure 4:** Features of *in silico* folded PUS7 mRNA targets. Related to Figure 4. (A) Metaplots of nucleotide frequencies for non-PUS7-dependent pseudouridine identified in (left) direct RNA UNUAR, (middle) MPRA *in vitro*, and (right) MPRA *in cellulo*. Position 0 represents the pseudouridine. For direct RNA, background set includes all U’s with adequate coverage. For the MPRA, background set includes any sequences not modified by PUS7 in the respective assay. (B) Minimum free energy was calculated for MPRA sequences using RNAfold. MPRA sequences not dependent on PUS7 *in vitro* or *in cellulo* with Nano-BID-Amp were compared to sequences found to be PUS7-dependent *in vitro* and *in cellulo* with Nano-BID-Amp. Mann-Whitney U test with Benjamini-Hochberg corrected p-value = 6.87 x 10^-14. (C) Metaplots of pairing probabilities calculated by RNAfold for non-PUS7-dependent pseudouridine identified in (left) direct RNA UNUAR, (middle) MPRA *in vitro* Nano-BID-Amp, and (right) MPRA *in cellulo* Nano-BID-Amp. Position 0 (gold dot) represents the pseudouridine. Background set excluded any sites found to be PUS7-dependent in respective assay. Direct RNA site list was down-sampled to only include 50 sites in UNUAR motif.

**Supplemental Figure 5:** Analysis of PUS7 sequence and structural features by modification quartile. Related to Figure 5. (A) Nucleotide frequency metaplots quartiles defined by average deletion rate in BS-treated samples, as defined in Figure 5A, for sites modified *in vitro* (left) or *in cellulo* (right) in Nano-BID-Amp MPRA. Position 0 represents the pseudouridine. (B) Minimum free energy (MFE, top) and maximum expected accuracy (MEA, bottom), determined with SHAPE-constrained RNAfold, for MPRA sites modified (left) *in vitro* or (right) *in cellulo* in Nano-BID-Amp, divided into quartiles as defined in Figure 5A. Mann-Whitney U test with Benjamini-Hochberg corrected p-value = 0.0072 for *in cellulo* MFE and p-value = 0.0120 for *in cellulo* MEA, between Q1 and Q4. (C) Average pairing probability, determined with SHAPE-constrained RNAfold, metaplots quartiles defined by average deletion rate in BS-treated samples, as defined in Figure 5A, for sites modified *in vitro* (top) and *in cellulo* (bottom) in Nano-BID-Amp MPRA. Position 0 represents the pseudouridine. Abbreviations: Q1, first quartile; Q2, second quartile; Q3, third quartile; Q4, fourth quartile; BS, bisulfite-treated Notes: * p < 0.05, ** p < 0.01

**Supplementary Figure 6:** Characterization of PUS7-dependent pseudouridines by cell type. Related to figure 6. (A) Out of 178 PUS7-dependent pseudouridine identified with Nano-BID-Amp *in cellulo* MPRA, 49 exhibited a higher deletion rate in HEK293T cells than HepG2 cells with a Δ deletion rate > 0.05 and Benjamini-Hochberg-corrected p-value < 0.05 (n = 3, multi-way ANOVA). (B) Heatmap of BS-treated deletion rates in WT *in cellulo* MPRA Nano-BID-Amp for 49 PUS7-dependent pseudouridine with a Δ deletion rate > 0.05 between HepG2 and HEK293T and a Benjamini-Hochberg-adjusted p-valued less than 0.05 (n = 3, multi-way ANOVA). (C) Immunofluorescence in HepG2 and HEK293T WT cells of DAPI and anti-PUS7 (abcam). Antibody specificity validated through KO (third row). PUS7 overexpression had similar localization patterns (fourth row). (D) Western blot (left) of seven different cell types (equal amounts loaded) RIPA lysates using Sigma PUS7 antibody. GAPDH is included as a loading control. Quantification with ImageJ (right) of PUS7 normalized to GAPDH. (E) Level of PUS7 mRNA in cancer cell types compared to normal tissue. Data was collected from the GEPIA web server (Tang, Z. et al. (2017) GEPIA: a web server for cancer and normal gene expression profiling and interactive analyses. Nucleic Acids Res, 10.1093/nar/gkx247). Three methods for differential expression analysis were used (ANOVA, LIMMA and Top 10) that consider RNA-seq data from TCGA tumor samples with paired adjacent TCGA normal samples and GTEx normal samples. Abbreviations: CHOL, Cholangio carcinoma; COAD, Colon adenocarcinoma; DLBC, Lymphoid Neoplasm Diffuse Large B-cell Lymphoma; ESCA, Esophageal carcinoma; GBM, Glioblastoma multiforme; LAML, Acute Myeloid Leukemia; LUSC, Lung squamous cell carcinoma; READ, Rectum adenocarcinoma; STAD, Stomach adenocarcinoma; TGCT, Testicular Germ Cell Tumors; THYM, Thymoma

## Methods

### Cloning

To generate overexpression vectors, PUS7 cDNA was cloned into a pLentiCMV backbone with a N-terminal V5 tag via Gibson assembly. For an empty control, the V5 tag alone was left in the pLentiCMV backbone. To generate constitutive PUS7 KD, shRNA against PUS7 was cloned into a pLKO backbone under a U6 promoter. pLKO.1 – TRC Cloning Vector (Addgene) was digested with EcoRI and AgeI to remove the stuffer sequence. Two oligos containing the complementary shRNA targeting sequence with the corresponding overhangs were annealed and ligated into the vector by standard cloning: shPUS7_4F 5’-CCGGTCTTAGTTCAGACTCATATATCTCGAGATATATGAGTCTGAACTAAGATTTTTG –3’, shPUS7_4R 5’-AATTCAAAAATCTTAGTTCAGACTCATATATCTCGAGATATATGAGTCTGAACTAAGA-3’. An empty vector control was generated by filling in the gaps with Klenow and blunt ended ligation.

### MPRA Generation

To build the MPRA (pool 1), PUS7-dependent pseudouridines from our Nanopore direct RNA sequencing and from precious studies^10,12,16^. Genomic sequences were extracted from the GRCh38 genome to center the psi in a 130-nucleotide sequence. To the 5’ end, a T7 promoter (GCTAATACGACTCACTATAG) and universal adapter (GACGCTCTTCCGATCT) were appended. To the 3’ end, a universal adapter (CACTCGGGCACCAAGGAC) was appended. The pool was ordered from Twist Bioscience.

To generate a vector for *in cellulo* transient transfection, PCR was performed on the pool using the universal handles and Phusion polymerase (New England Biolabs #M0530L) for 10 cycles with primers for overlap with the backbone. A SFFV-mCherry backbone, a kind gift from Xiaojing Gao, was digested with MluI and ApaI to create an opening in the 3’ UTR of mCherry. Gibson assembly with HiFi Assembly mix (New England Biolabs #E2621L) was performed. The resultant product was purified with Zymo DNA Clean and Concentrator-5 (Zymo Research #D4014) and eluted into 10 ul water. Endura electrocompetent cells (Biosearch Technologies #60242-1) were thawed on ice, 25 ul of cells were combined with 2 ul of purified vector. Enough bacteria for 1000x coverage of all sequences were transferred into 0.2 cm cuvettes and electroporated on a Biorad electroporator with settings of 10 uF, 600 Ohms, and 1800 Volts. Recovery media was immediately added to cells per manufacturer’s instructions. Cells were grown at 200 rpm and 37C for 1 hour before being pooled and plated equally across 4 large LB + ampicillin bioassay plates. Cells were grown overnight. Then next day, all colonies were collected and plasmid DNA was extracted with two columns from PureLink HiPure Plasmid Filter Midiprep Kit (Thermo Fisher # K210014). To confirm pool presence and distribution, the same MPRA primers used in Nano-BID-Amp were used to amplify off the plasmid. The plasmid was sequenced in accordance with Nanopore cDNA Ligation kit (SQK-LSK114) and PCR Barcoding kit (EXP-PBC096) protocols.

### Cell Culture Standards

Human hepatocellular carcinoma cells HepG2 from ATCC HB8065 (lot 59635738) and human embryonic kidney cells HEK293T (kind gift from Dr. Lingyin Li) were grown in high glucose DMEM (ThermoFisher #11965118) supplemented with 10% fetal bovine serum (FBS HyClone SH30071.03) and 1% Penicillin-Streptomycin (ThermoFisher #15140163). Cells were grown at 37C with 5% CO2 and maintained at sub-confluency.

### Cell Line Creation

In this work, we generated cells lines for PUS7 overexpression, PUS7 dox-inducible knockdown, and PUS7 constitutive knockdown in HepG2 and HEK293T cells with a lentiviral approach. First, lentivirus was generated with Lipofectamine 2000 (Thermo Fisher #11668030) wherein 2 ug of desired vector was mixed with 1.8 ug pCMV-psPAX2 and 0.2 ug pCMV-VSV-G according to manufacturer’s instructions. Solution was plated dropwise onto adherent HEK293T cells in a P6 dish at 70% confluency. Three days later, media was filtered through a 0.45 um cellulose acetate filter to collect viral particles. HEK293T and HepG2 cells had been plated in a 6 well plate at 100,000 cells per well the day prior. Media was removed from these cells and replaced with 1 ml of fresh media, 1 ml of virus, and 1.6 ul of 10 mg/ml polybrene solution. Media was changed 24 hours later and cells left to grow undisturbed for a total of 72 hours. At that point, cells were placed under 3 ug/ml puromycin or blasticidin selection (as appropriate) for 48 hours, until an uninfected control perished. Cells were allowed to recover for over 48 hours before a sample was collected for PUS7 expression assessment by Western blot.

### Dox-Inducible PUS7 shRNA knockdown

HepG2 or HEK293T dox-inducible cells were plated at subconfluency with a working concentration of 2 ug/ul doxycycline (Sigma Aldrich # D5207-10G). 48 hours later, cells were expanded and fresh doxycycline at 2 ug/ul were added. Cells were collected 96 hours after initial administration of doxycycline. To collect, media was aspirated from plate. Trizol (ThermoFisher #15596018) was added directly to the plate, per manufacturer’s instructions. Cells were homogenized in Trizol and processed immediately or stored at –80C until needed. Western blot of PUS7 depleted lysates confirmed depletion to ∼90%.

### Nanopore direct RNA sequencing

Total RNA was isolated from stable HepG2 cell lines expressing empty vector pLKO-Tet-On or vector with a Dox-inducible shRNA targeting PUS7: shPUS7_4F 5’-CCGGTCTTAGTTCAGACTCATATATCTCGAGATATATGAGTCTGAACTAAGATTTTTG-3’ shPUS7_4R 5’-AATTCAAAAATCTTAGTTCAGACTCATATATCTCGAGATATATGAGTCTGAACTAAGA-3’ from 3 biological replicates as described previously^10^. shRNA expression was induced in the stable HepG2 cell lines with Doxycycline to a final concentration of 500 ng/mL. Cells were maintained in Doxycycline containing media for 96h. Nanopore direct RNA sequencing was carried out with kit SQK-RNA004 and each sample was sequenced on an individual FLO-MIN004RA or FLO-PRO004RA flow cell to sequencing depths of 2-6M reads. Sequencing of the synthetic UNΨAR RNA Concatemer standard was carried out by barcoding each mix of defined ratios on modified to unmodified from 0-100% and demultiplexed using SeqTagger^62^.

### Nanopore direct RNA sequencing Data Analysis

Multi-pod5 files from three replicates of PUS7 WT/KD Nanopore direct RNA sequencing data were basecalled with Dorado (7.3) using the high accuracy model. Reads were subsequently aligned to human genome assembly GRCh38 from GENCODE using minimap2 with the option “-ax splice –k14”. Resultant SAM files were converted to BAM files using Samtools (1.16.1) and subsequently visualized in IGV.

### PUS7-Dependent Pseudouridine Site Assignment

A custom script, ModDetect, was developed based on PsiDetect^34^ to call putative RNA modification sites from mutational signatures in sequencing data from a test sample compared to a control. Extending PsiDetect’s original function of detecting pseudouridines from direct RNA U-to-C mismatches, ModDetect also enables readouts of pseudouridines based on deletion signatures in bisulfite-treated samples as well as detection of any other modified bases that manifest as mutations in sequencing reads. ModDetect was used to called PUS7-dependent pseudouridine sites from direct RNA U-to-C mismatches. We filtered out homopolymers as they are known to be error prone in Nanopore direct RNA sequencing^63^. The following criteria were used to call PUS7-dependent pseudouridine sites: (1) number of T+C reads >= 10 in both WT and KD libraries; (2) U-to-C mismatch >= 15% where U-to-C mismatches rates were calculated as C/(T+C); (3) site is called in at least 2 out of 3 replicates. To account for sites that might have been missed due to insufficient coverage in our libraries, an additional set of sites was called which met criteria (1) and (2) in at least 1 of our own replicates and was also annotated as a PUS7-dependent pseudouridine site in an external dataset^10,12,16^.

### UNΨAR RNA Concatemer Design

To derive pseudouridine stoichiometries in the consensus PUS7 UNUAR motif, a synthetic UNΨAR RNA concatemer was ordered from TriLink. It was designed to contain the 8 possible UNΨAR sequences separated by 14-nt spacer sequences, with a 10-nt poly-A tract placed at the 3’ end for compatibility with the standard Nanopore direct RNA sequencing kit SQK-RNA004. The spacer sequences were identical to those in a previous synthetic RNA oligo library^12^.

### UNΨAR direct RNA Stoichiometry Derivation

To make calibration curves, data from each motif was independently plotted, with the background U-to-C mismatch rate at 0% UNΨAR subtracted from higher mixes. A linear line of best fit was applied to each curve and the equation was extracted (Table S2). To correct PUS7 site mismatch rate to real stoichiometry, the averaged mismatch rate for a given site across the replicates was used in the equation appropriate for the motif of the site.

### Transient Transfection for In Cellulo MPRA

Prior to primary experiment, pilots were performed to determine transfection efficiency. The determined transfection efficiency was used to calculate how many cells to plate to ensure 1000x coverage of all sequences in the pool (cells needed = (# sequences in pool x 1000) / transfection efficiency). Cells were plated 24 hours prior, targeting 70% confluency with at least the cell number calculated by time of transfection.

Manufacturer’s instructions for Lipofectamine 3000 (ThermoFisher # L3000008), scaled to the appropriate size, were followed for transfection. Cells were incubated for 48 hours without media change. Cells were visualized under fluorescent microscope to establish transfection efficiency. To collect, media was aspirated from plate.

Trizol (ThermoFisher #15596018) was added directly to the plate, per manufacturer’s instructions. Cells were homogenized in Trizol and processed immediately or stored at –80C until needed.

### RNA Extraction and polyA Selection

RNA was extracted following manufacturer’s instructions for Trizol (ThermoFisher #15596018). In brief, chloroform was added to homogenized cell-Trizol mix, mixed thoroughly, and incubated at room temperature for 3 minutes. Samples were centrifuged at 12,000g for 15 min at 4C. The upper aqueous phase was transferred to a new tube for RNA extraction. The lower organic phase was transferred to a new tube for protein extraction. An equal volume of isopropanol was added. Samples were incubated on ice for 10 min.

Samples were centrifuged at 12,000g for 10 min at 4C. Supernatant was discarded. RNA pellet was washed once with 75% ethanol and centrifuged at 7,500g for 5 min at 4C. Supernatant was discarded. Pellet was air dried and resuspended in RNAse free water. Concentration was quantified by Qubit RNA Broad Range Assay (ThermoFisher #Q10211).

Two rounds of polyA selection were performed with Oligo d(T) Magnetic Beads (New England Biolabs #S1419S) on ∼35 ug of total RNA. In brief, beads were equilibrated in Binding Buffer. RNA was denatured at 65C for 2 min and immediately placed on ice. RNA was added to beads and rotated at room temperature for 5 min. Samples were placed on magnets and washed twice with Washing Buffer B. Following final wash, all supernatant was removed. Cold 10 mM Tris HCl was added to samples. Samples were incubated at 75C for 2 min, placed on magnet, and quickly transferred to a new RNAse-free tube. Concentration was quantified by Qubit RNA High Sensitivity Assay (ThermoFisher #Q32855).

### Protein Extraction

From the Trizol extraction, 150 ul of the lower organic phase was combined with 1350 ul of ice-cold methanol. Samples were vortexed to mix. Samples were centrifuged at 14,000g for 10 min at 4C. Supernatant was discarded. Samples were washed with 1 ml of ice-cold methanol. Samples were centrifuged at 14,000g for 10 min at 4C. Supernatant was discarded. Samples were washed with 80% ethanol. Samples were spin at 14,000g for 10 min at 4C. Supernatant was discarded. Protein pellet was air-dried at room temperature.

Samples were resuspended in 1% SDS 100 mM TAEB, freshly made. Samples were incubated at room temperature overnight for full resuspension. Samples were quantified via BCA assay (ThermoFisher #23227) or Qubit Protein Broad Range Assay (ThermoFisher #A50669). Samples were stored at –20C until needed.

### In Vitro Transcription

In vitro transcription reactions were performed according to the Megashort script Kit (ThermoFisher AM1354). Briefly, 10X T7 reaction buffer, and 4 nucleotide triphosphate solutions (75mM each) were thawed at room temperature, briefly vortexed, and spun down. Each transcription reaction contained 2µL of 10X T7 reaction buffer, 2 µL of each NTP solution, 8µL of template DNA (2 picomoles) and 2 µL of T7 Enzyme mix. In vitro transcription reactions were incubated 37°C for 4 hours followed by the addition of 2 units of Turbo DNAse. An equal volume of gel loading buffer (95% Formamide, 18 mM EDTA, and 0.025%, Xylene Cyanol, and Bromophenol Blue) was added and full-length RNA molecules were resolved and extracted from 8% polyacrylamide, 7M Urea gels under blue light after 5 minutes of staining with SYBR Gold. RNA was eluted with 1mL 300mM NaOAC, 1mM EDTA supplemented with RNAse inhibitor overnight at 4°C. RNA eluates were passed through a Corning SpinX column spun at 10,000xg for 5 minutes followed by the addition of an equal volume of isopropanol and 1.5 µL 20 mg/mL glycogen. RNA precipitation reactions were incubated at –20°C overnight and spun 21,300xg at 4°C for 20 minutes. Precipitated RNA pellets were washed 2 times with 70% ethanol and resuspended in 100 µL nuclease free H2O per pellet. RNA was quantified using the Qubit RNA BR assay kit.

### PUS7 protein purification

BL21(DE3) Gold competent cells (Agilent #230132, USA) were transformed with 50 ng of pET15b PUS7-His plasmid. Recombinant protein induction was performed with 0.1 mM IPTG (Sigma Aldritch # I6758-5G, USA) at OD600 1-1.2 for 16 hours at 20°C in 6 L of TB medium. Cells were harvested by centrifugation at 4000g and 4°C for 45 minutes and bacterial pellets were resuspended in lysis buffer (50 mM potassium phosphate buffer pH 8, 200 mM NaCl, 0.1% Triton X-100, 1 mM DTT, 1x Halt Protease Inhibitor Cocktail [Thermo Fisher #78430,USA], 1 mM PMSF, 10U/mL benzonase [Millipore # E1014, USA] and 1mg/mL lysozyme [Thermo Fisher #89833, USA]) and sonicated 4 times using an amplitude of 60% for 30 seconds followed by 1 minute incubation on ice. Lysates were spun down at 12000g for 30 minutes at 4°C and the supernatant was filtered through a 0.2 μm PES filter (Thermo Fisher #725-2520, USA). Cleared lysate was used for metal affinity chromatography using an ÄKTA FPLC system (Cytiva, USA) using a HiTrap IMAC Sepharose FF 5mL column previously equilibrated with 2 column volumes of lysis buffer. After sample application column was washed with 4 column volumes of wash buffer (50 mM potassium phosphate buffer pH 8, 500 mM NaCl, 30 mM Imidazole, 1 mM DTT, 1 mM PMSF). Protein was eluted using a gradient of 0-95% elution buffer (50 mM potassium phosphate buffer pH 8, 500 mM NaCl, 300 mM Imidazole, 1mM DTT, 1 mM PMSF) in 1 mL fractions. Fractions with the highest protein content were pooled and used for buffer exchange with a 10 mL Zeba spin desalting column (Thermo Fisher #89894, USA) into storage buffer (20 mM HEPES (pH 7.5), 200 mM NaCl, 10% glycerol, 1 mM DTT, 1x Halt Protease Inhibitor Cocktail [Thermo Fisher, #89833USA], 1mM PMSF).

Recombinant protein was concentrated using Amicon ultra centrifugal filter 10 kDa MWCO (Millipore, USA) for 7 minutes at 4000g and 4°C in a swinging bucket rotor and aliquots were flash frozen in liquid nitrogen. Protein concentration was determined by BCA (Thermo Fisher, USA#23227) using BSA standards and a NanoDdrop One spectrophotometer (Thermo Fisher, USA).

### In Vitro Pseudouridylation

In vitro pseudouridylation reactions were performed as previously described^10,28,39^. Briefly, 30 picomoles of purified oligo pool was diluted into 39.5 µL nuclease free H2O per pseudouridylation reaction. RNA was heated to 75°C for 2 minutes and then snap cooled on ice for 5 minutes. Refolding was initiated with the addition of 10 µL 5X pseudouridylation buffer (500 mM Tris-HCl pH 8.00, 500mM NH4OAC, 25mM MgCl2, and 1.5mM EDT). RNA was incubated at 37°C for 20 minutes. In vitro pseudouridylation reactions were prepared by combining 90 µL 5X pseudouridylation buffer, 10 µL 100mM DTT, recombinant PUS7 to 600nM, 46.5 µL refolded RNA, and H2O to 500 µL. Pseudouridylation reactions were incubated at 30°C for 45 minutes and then extracted with acid phenol:chloroform and precipitated with isopropanol as previously described.

### Nanopore and Illumina Sequencing Comparison

Synthetic 4 uridine-containing standard (4U) was purchased from IDT (USA) with the following DNA sequence (uridines are bolded): GGAACAGAAACAGAGAAAGGAACAGAGAAAGACA**T**AAACAGAAAGAGACAAGAACAGAGACAAGAACAG**T** GGCAGGAACAGAGACAAACAGAGACAGGAACAA**T**GACAGGAACAGAAAGAAACAGAGACAAGCAC**T**CGG GCACCAAGGACACGAACCGGAACGCGGAACCAAACGGGCAACGGACCGGAC. The DNA was used for In vitro transcription reactions using 200 nM of the synthetic template in the presence of either UTP or ΨTP using the T7 MEGAshortscript in vitro transcription kit (Thermo Fisher, #AM1354 USA). Reactions were TURBO DNase-treated (Thermo Fisher, #AM1907 USA) and gel purified through denaturing Urea PAGE. Gel slices were eluted overnight in RNA elution buffer (300 mM sodium acetate pH 5.3, 1 mM EDTA pH 8, supplemented with 100 U/mL RNAse inhibitor (Promega, USA) # N2615) at 4°C. RNA was precipitated in isopropanol, resuspended in RNAse-free water and quantified using a Qubit 4 fluorometer (Thermo Fisher, USA). Twenty nanograms of either 4U or 4Ψ were used for bisulfite treatment with previously described conditions.

Desulfonated eluted RNA was used for reverse transcription reactions using Superscript IV (Thermo Fisher #18090200, USA) as previously described and using a template-specific reverse primer (5’ GTCCGGTCCGTTGCCCG 3’). Reactions were further incubated with 5 units of RNase H (New England Biolabs # M0297L, USA) for 20 minutes at 37°C followed by 5 minutes at 70°C and cDNA was purified using Zymo DNA Clean & Concentrator Kit (Zymo Research # D4014, USA). The cDNA was used for PCR reactions using the primers fw: 5’ TTTCTGTTGGTGCTGATATTGCGGAACAGAAACAGAGAAAGGAACAG 3’ and rev: 5’ ACTTGCCTGTCGCTCTATCTTCGTCCGGTCCGTTGCCCG 3’ and Phusion High-Fidelity DNA Polymerase (New England Biolabs # M0530L, USA). Products were purified using Zymo DNA Clean & Concentrator Kit (Zymo Research # D4014), USA) and used either for Nanopore sequencing using the ligation sequencing DNA V14 kit SQK-LSK114 (Oxford Nanopore, United Kingdom) or submitted for Illumina sequencing (Genewiz Azenta, USA). Reads were mapped using Minimap2 ^64^.

### Nano-BID-Amp

Twenty nanograms of RNA from in vitro pseudouridylation or double-polyA selection was prepared in 10.5 ul nuclease-free water in PCR strip tubes. Immediately before beginning the reaction, a bisulfite mix of 2.4 M Na2SO3 and 0.36 M NaHSO3 was resuspended in 1 ml of nuclease free water preheated to 70C. RNA samples were transferred to room temperature and 45 ul of bisulfite mix was added to each reaction well.

Samples were incubated at 70C for 3 hr with gentle shaking. Zymo RNA Clean and Concentrator 5 kit (Zymo Research #R1016) was used to purify RNA as follows: samples were combined with 75 ul water, 270 ul RNA Binding Buffer (Zymo Research #R1013-2-100), 400 ul 100% ethanol, mixed thoroughly, loaded on the column and spun as instructed in kit manual. Samples were washed once with 200 ul RNA Wash Buffer (Zymo Research #R1003-3-48) as instructed. 200 ul of RNA Desulphonation Buffer (Zymo Research # R5001-3-40) was added directly to the column, samples were incubated at room temperature for 1 hr 15 min. Columns were spun, desulphonation buffer was discarded. Samples were washed twice more with RNA Wash Buffer (700 ul for first wash, 400 ul for second wash) until completely dry. Samples were eluted in 11 ul nuclease free water and immediately proceeded to reverse transcription.

For reverse transcription, bisulfite-treated RNA was prepared in 10.5 ul nuclease-free water with a matched untreated input control of 20 ng RNA in 10.5 ul water. We found that bisulfite treatment degraded approximately 50% of RNA, therefore, we recommend performing two bisulfite reactions (combined before desulphonation) for every one input reaction, especially for low abundance transcripts. One microliter of 2 uM template-specific primers with a 10mer UMI and Nanopore adapters handles were added to the reaction. For MPRA reactions, samples were incubated at 65C for 2 min. For endogenous transcript reactions, samples underwent a touchdown annealing wherein temperature started at 90C and declined 2C per minute until reaching 50C. Samples were held at 50C and 8.5 ul of RT master mix (4 ul 5X SuperScript IV Buffer, 2 ul 10 mM dNTPs, 1 ul 100 mM DTT, 0.5 ul RNasin Plus, and 1 ul SuperScript IV (Thermo Fisher #18090200)) was added to each reaction. Samples were incubated at 50C for 1 hr. 2 ul of 4M NaOH was added to degrade RNA, samples were incubated at 95C for 3 min before proceeding immediately to bead clean-up. A 1x volume of RNAClean XP beads (Beckman Coulter #A63987) were used in a single clean-up step to remove primers. Samples were eluted into 20 ul cDNA.

For the first round of PCR, a 20 ul reaction was set up with 4ul of eluted cDNA, 10 uM of primers (containing Nanopore adapter sequences), and Phusion polymerase (New England Biolabs #M0530L). PCR was optimized to the specific primers used, with between 20 and 35 cycles. PCR success was assessed with TapeStation D1000 ScreenTape (Agilent #5067-5582) to check product size and abundance. DNA was cleaned with AMPure XP beads (Beckman Coulter #A63881), using a 1.8x bead ratio, per manufacturer’s instructions. DNA was eluted in up to 20 ul of nuclease-free water. Samples were then proceeded through the Nanopore cDNA Ligation (Oxford Nanopore Technologies #SQK-LSK114) with PCR Barcoding Expansion (Oxford Nanopore Technologies #EXP-PBC096) kit per manufacturer’s instructions. Barcoded samples were pooled and sequenced on either a Flongle (R10.4.1) or Promethion (R10.4.1) flow cell.

### Nano-BID-Amp Bioinformatic Analysis

Nanopore sequencing produced several fastq files for each barcode. The fastq files were concatenated together for each barcode and renamed based on the sample name. Nanopore adapters were trimmed using cutadapt and UMIs extracted with UMI tools. For MPRA experiments, universal handles were trimmed with cutadapt. Samples were mapped with Minimap2 using parameters –k5. Reads were deduplicated with UMI tools. Files were converted to bam, sorted, and indexed. A custom Rscript used package pileup to count the nucleotide calls at each position. Samples were filtered to only contain positions with at least 20 reads of coverage. Deletion rates were calculated as (deletion counts / total reads).

To determine the effect of bisulfite treatment on deletion rate, first samples are filtered to contain only those with a greater than 0.05 difference in deletion rate between the average deletion rate in bisulfite-treated samples (wild-type only for in cellulo) and input samples. Next, a generalized linear model (glm) was built to measure the impact of random effects (replicate, etc) and treatment (fixed effect) on the deletion rate (response). The model is weighted so samples with higher total read count carry more weight, as a higher read depth (sample size) reduces the impacts of statistical noise. An ANOVA test is then run on this model to determine significance. Multiple-hypothesis test correction is performed with a Benjamini-Hochberg calculation. The corrected p-value is then filtered for p < 0.05; all sites that fall below this threshold are called as significant.

To determine the impact of tertiary factors (PUS7 expression, cell type) on deletion rate, a similar process was followed. First, first samples are filtered to contain only those with a greater than 0.05 difference in deletion rate between the average deletion rate in the two factors under consideration (WT and KD, WT and OE, HepG2 and HEK293T). Next, two generalized linear models (glm) are fit to the data, one accounting for the factor (PUS7 level, cell type) having an effect and the other not. In both models, the deletion rate is the response, the factor is a fixed effect, the replicate is a random effect, and each sample is weighted by its total reads. The factor model has two indicators, L1andBS.ind and L2andBS.ind. L1 and L2 represent the two levels of the factor (WT and KD, WT and OE, HepG2 and HEK293T). These indicators carry a value of 1 if the sample is positive for both the level and BS treatment (so WT + BS for L1andBS.ind) and 0 in all other cases. The no factor model has one indicator, BS.ind, that is 1 is the sample is positive for BS and 0 if not. All three indicators are treated as fixed effects in their respective models. An ANOVA test then asks which model better fits the data for the site: distinguishing the factor levels or not. This is based on the logic that if there is no meaningful difference between the deletion rates in the different conditions, either model would fit equally well. However, if there was a meaningful difference between the levels, the factor-aware model would fit better, producing a significant p-value. All p-values then undergo multiple hypothesis test correction via Benjamini-Hochberg. The corrected p-value is then filtered for p < 0.05; all sites that fall below this threshold are called as significant.

The treatment analysis and PUS7 expression level analysis was performed on HepG2 and HEK293T data considered together, HepG2 data considered singularly, and HEK293T data considered singularly. The treatment analysis for *in cellulo* Nano-BID-Amp produced a list of sites that could be modified by any PUS. To determine PUS7-dependency in the in cellulo MPRA, the union of sites sensitive to PUS7 KD or PUS7 OE across any cellular background was taken. To determine PUS7-dependency in the *in vitro* MPRA, the site list from the treatment analysis was used as no other PUS was present.

### In Vitro Nano-SHAPE-Amp

In vitro SHAPE reactions were performed as described by Spitale and colleagues and MaP reverse transcription reactions were performed as previously described^45^. Briefly, 500 ng of oligo pool MPRA RNA not treated with recombinant PUS7 was diluted into 18 µL H2O. Samples were incubated at 95°C for 2 minutes and then snap cooled on ice for 2 minutes followed by the addition of 9 µL 3.3X RNA folding buffer (333 mM HEPES, pH 8.0, 20 mM MgCl2 and 333 mM NaCl). RNA folding was allowed to proceed at 37°C for 10 minutes and then 3 µL 1M NAI (2-methylnicotinic acid imidazolide) or 100% DMSO was added. SHAPE reactions were stopped after 15 minutes by the addition of 170 H2O and 200 H2O Acid Phenol:Chloroform. Samples were vortexed, spun at 21,300 xg for 5 minutes and the aqueous phase was moved to a clean tube. Residual phenol was removed by the addition of an equal volume of chloroform followed by a vortex, spin at 21,300 x g and removal of the aqueous phase to a clean tube. RNA was precipitated with addition of .1 volume of 3M NaOAC, 2 µL 20 mg/mL glycogen and 3 volumes of 100% ethanol. Samples were incubated at –80°C for 30 minutes and then centrifuged at 21,300 x g, 4°C for 30 minutes. RNA pellets were washed twice with 70% ethanol and centrifuged at 21,300 x g 4°C for 15 minutes after each wash. Each RNA pellet was resuspended in 8.5 µL H2O. For primer annealing, 2 µL 10 µM RT primer and 1 µL of 10mM dNTPs were added to each reverse transcription (RT) reaction. RT Primers were annealed by incubating reactions at 70°C for 5 minutes followed by snap cooling on ice. After annealing, 4 µL 5X RT buffer (250 mM Tris HCl pH 8.0, 375 mM KCl) 2 µL .1M DTT, .5 µL Murine RNAse inhibitor (New England Biolabs # M0314L) and 1 µL 120 mM MnCl2 were added to each reaction. RT reactions were brought up to 42 °C and then 1 µL Superscript II RT (Thermo Scientific #18064014) was added to each reaction and mixed. Reverse transcription was performed at 42°C for 2 hours, 50°C for 10 minutes and 55°C for 10 minutes. To remove RNA post RT, 2 µL of 4M NaOH was added and reactions were incubated at 95°C for 3 minutes. The volume of RT reactions was adjusted to 50 µL and cDNAs were purified twice with a 1.8X volume of AMPure XP beads (Beckman Coulter # A63881) as previously described to remove UMI containing RT primers. Nanopore amplicon sequencing libraries were created according to the Nano-BID-Amp library construction protocol.

### Nano-SHAPE-Amp Bioinformatic Analysis

Raw pod5 files from all samples were base-called using dorado 1.1.0 in super high accuracy mode (sup) with –-no-trim flag. Resultant bam files were demultiplexed with dorado demux with the –-no-trim and –-no-classify flags. Nanopore adapters were trimmed using cutadapt using linked adapters. To account for Nanopore sequencing occurring in the sense and antisense direction, this first adapter trimming was performed with sense and antisense primers. Reads matching the antisense primers were reverse complemented. Next, UMIs were extracted with UMI tools. Universal handles were trimmed with cutadapt using linked adapters. Samples were mapped with bowtie2 using parameters –-local –-sensitive-local –-ignore-quals –-no-unal –-mp 3,1 –-rdg 5,1 –-rfg 5,1 –-dpad 30. Reads were deduplicated with UMI tools and resultant bam files were converted to fastq files.

Samples were then analyzed with shapemapper2-2.3^46^ with parameters –-min-depth 1000 –-window-to-trim 10 –-min-qual-to-count 22 –-min-qual-to-trim 5 –-indiv-norm. SHAPE-treated samples were compared to DMSO-treated samples. Sequences lacking adequate quality, as indicated by shapemapper2-2.3, were filtered from the analysis. SHAPE reactivities were extracted for each sequence and averaged across replicates (if available) using inverse variance weighting.

### RNA Structure Analysis

RNA structure was determined with Vienna RNAfold^65^, with or without SHAPE-informed folding with command RNAfold –p [--shape=“$shape_file”] –-MEA. Minimum free energy (MFE) and maximum expected accuracy (MEA) values were extracted from each sequence. To create metaplots, pairing probabilities were extracted from RNAfold dp.ps output and averaged across all sequences considered. RNA structures were visualized using VARNA^66^ with the MEA dot-bracket notation.

### Structure Motif Description

To define the features of the Pupup signature, we created separate SHAPE-informed metaplots of PUS7 target sites identified in direct RNA sequencing, *in vitro* Nano-BID-Amp MPRA, and *in cellulo* Nano-BID-Amp MPRA. We generated a synthetic background set “random U” by randomly selecting 800 non-centered uridines from the Pool 1 MPRA located at least 30 nucleotides from the beginning or end of the target sequence. We re-centered the pairing probability of these random Us at position 0. To identify sites with statistically significant (p < 0.05) differences between PUS7 targets and the random U background, we performed Welch’s t-test with Benjamini-Hochberg correction on all sites +/-15 nucleotides from position 0. All statistically significant sites were included in the Pupup feature list.

### Immunofluorescence

Coverslips were placed into wells of a 24-well plate and sterilized with UV for 10 min, followed by a quick wash with DMEM. HepG2 cells were plated at a density of 100,000 cells per well; HEK293T cells were plated at a density of 50,000 cells per well. Cells were incubated overnight for attachment. Media was removed, cells were quickly washed with PBS. Cells were incubated with rocking in freshly diluted 4% paraformaldehyde in PBS for 10 minutes, followed by two washes with PBS. Cells were permeabilized with freshly diluted 0.5% Triton X-100 in PBS for 10 minutes with rocking and washed three times with PBS. Cells were blocked with IF blocking buffer (5% FBS in PBS) for 1 hour in a humidity chamber and 37C. Blocking buffer was removed and replaced with primary antibody (PUS7, abcam #ab289857) at a 1:500 concentration in IF blocking buffer for 1.5 hours.

Samples were washed with 1x Wash Buffer A (Millipore Sigma #DUO82049) two times for 5 minutes each. Samples with incubated with green anti-rabbit secondary (Invitrogen #A11008CATNO) at 1:1000 concentration in IF blocking buffer for 30 minutes, covered to protect from light. Samples were washed with 1x Wash Buffer B (Millipore Sigma #DUO82049) two times for 10 minutes each, followed by a final wash with 0.01x Wash Buffer B. Coverslips were mounted onto slides with In Situ Mounting Media with DAPI (Sigma-Aldrich #DUO82040), cured overnight, and imaged on ECHO Revolve microscope.

### Western Blot

RIPA lysis was performed on cell pellets, wherein cells were resuspended in RIPA 150 mM NaCl, 0.8% NP-40, 0.5% deoxycholate, 0.1% SDS, 5 mM Tris pH 7.4), incubated on ice for 10 minutes with periodic vortexing, and spun at max speed for 15 minutes at 4C. Supernatant was transferred to a new tube and quantified with Qubit Protein Broad Range Assay (ThermoFisher #A50669). Appropriate amounts of sample were loaded into a Precast Mini-PROTEAN TGX Gel (Biorad #4561096) and run at 130V for 50 minutes. Samples then underwent a wet transfer to PVDF membrane and were blocked for at least 10 minutes in 5% milk in TBST. Samples were incubated overnight in primary antibody at various concentrations in 5% milk in TBST. Samples were washed three times with TBST and incubated in secondary antibody at 1:4000 dilution for 1 hour. Following three washes, samples were imaged on a ChemiDoc MP Imaging System. Results were quantified in ImageJ and normalized to GAPDH loading control.

## Acknowledgements

We thank member of the Martinez Lab for feedback on the project and the manuscript. We thank generous colleagues who took the time to give us feedback on the manuscript. R00 GM135537, Rita Allen Foundation Award, Packard Fellowship for Science and Engineering from the David and Lucile Packard Foundation and Chan Zuckerberg Biohub San Fransisco Award to N.M.M. T32GM136631 from N.I.H. to R.R. and N.R.. VPUE Developmental Biology Undergraduate Research Grant to R.J.

Research reported in this publication was supported by the National Institute Of General Medical Sciences of the National Institutes of Health. The content is solely the responsibility of the authors and does not necessarily represent the official views of the National Institutes of Health.

