## Supplemental Figures for "Nano-Mod-Amp reveals RNA sequence, structural and cell type specific features of pseudouridylation by PUS7"

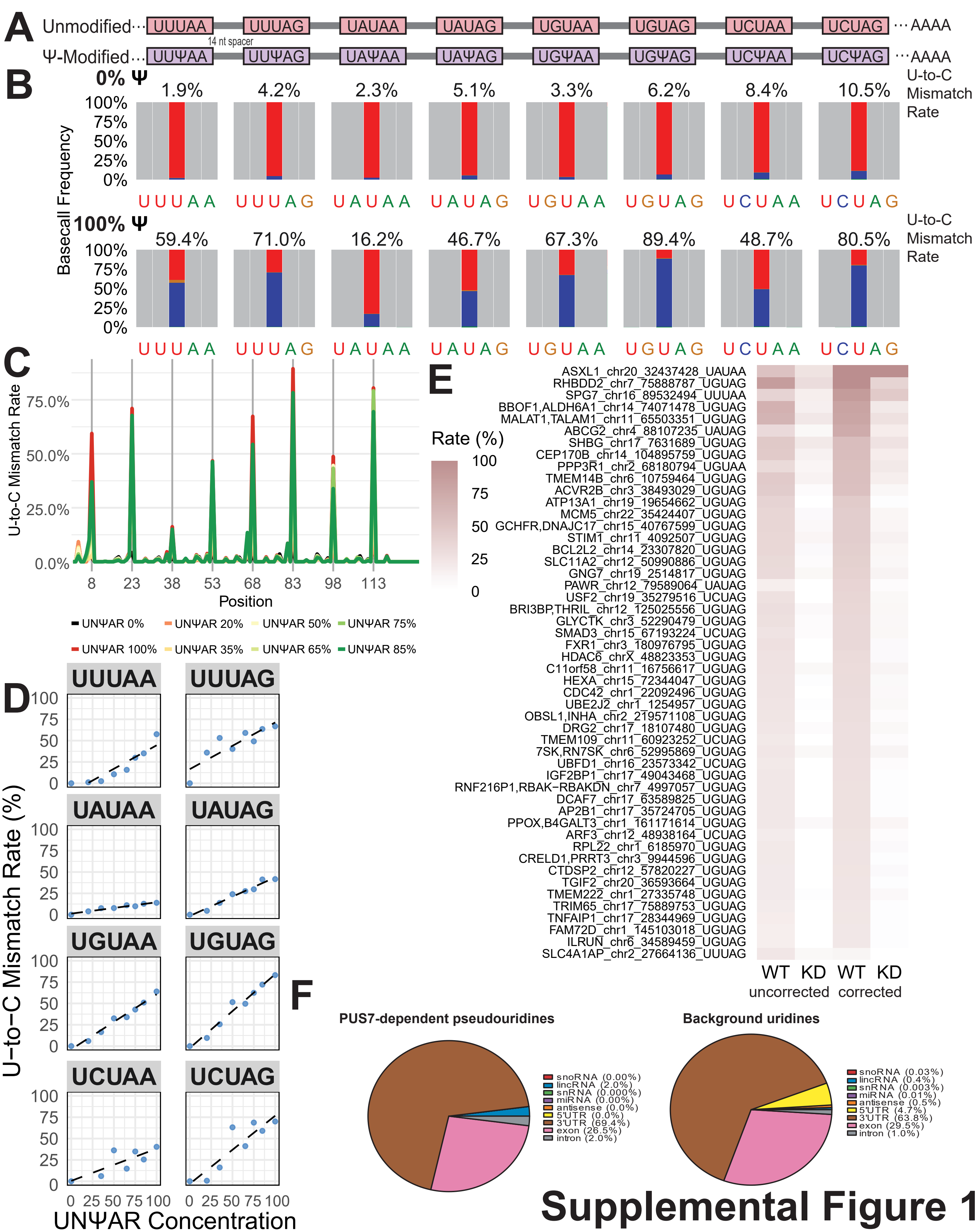


### Supplemental Figure 1

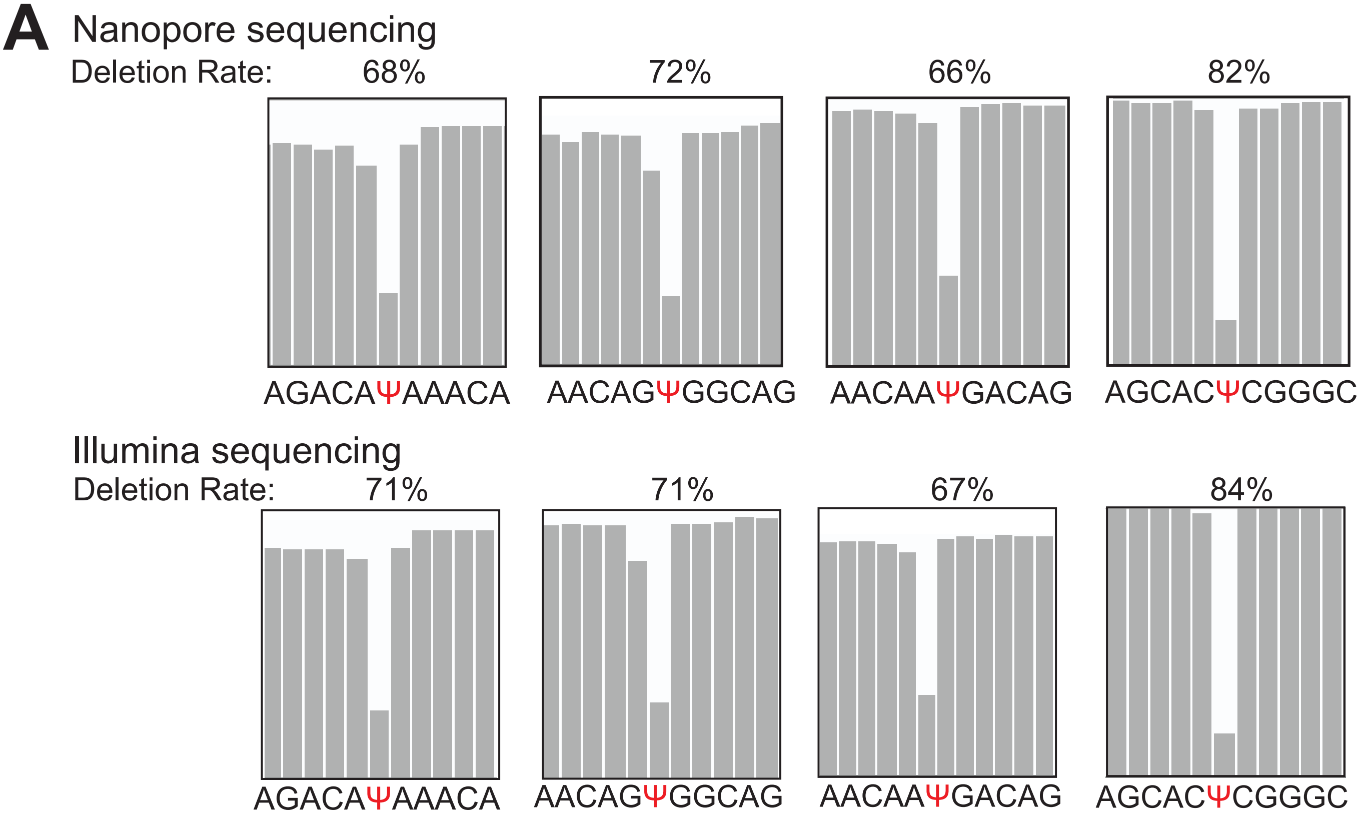


**Supplemental Figure 2**

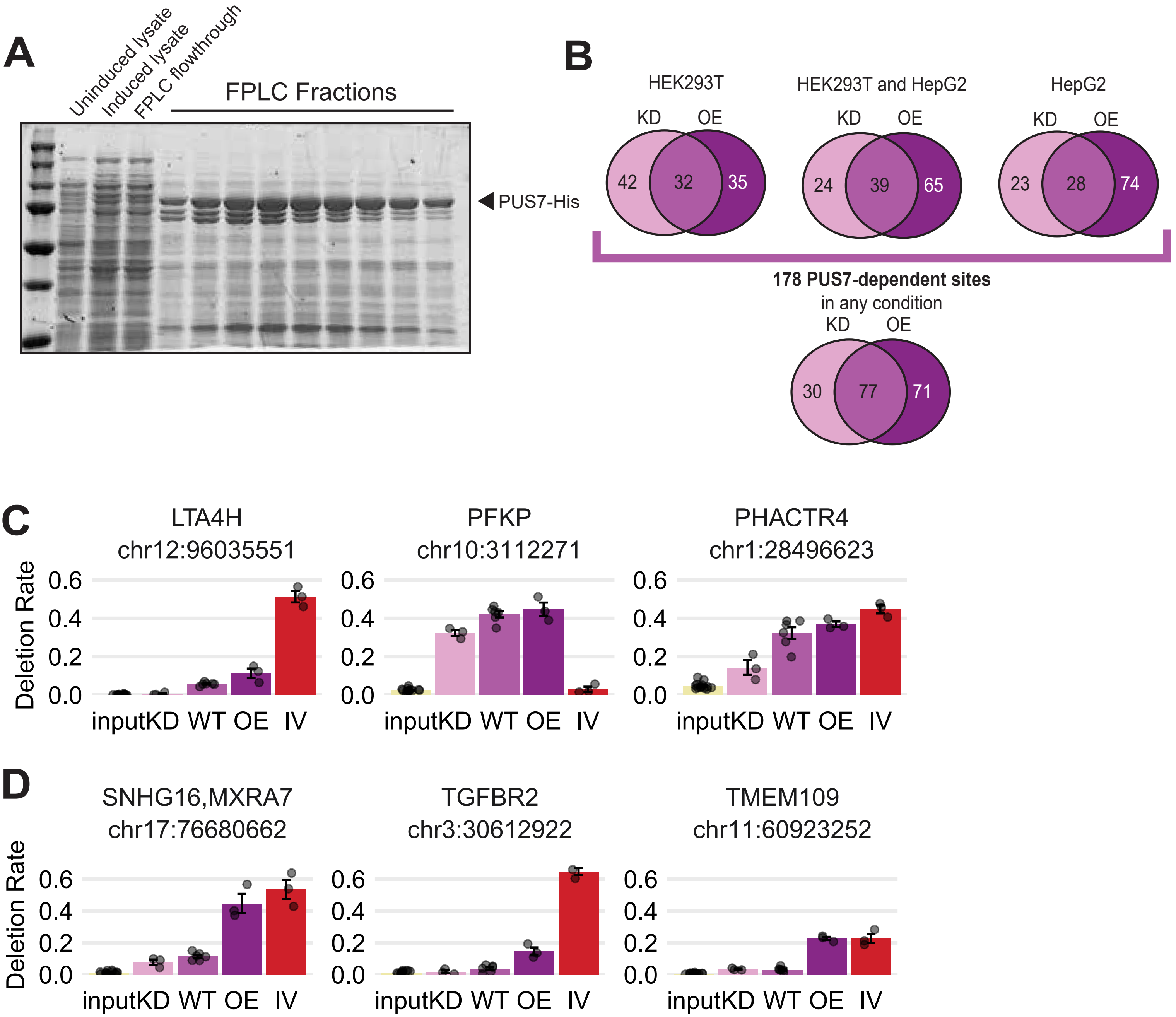


Supplemental Figure 3

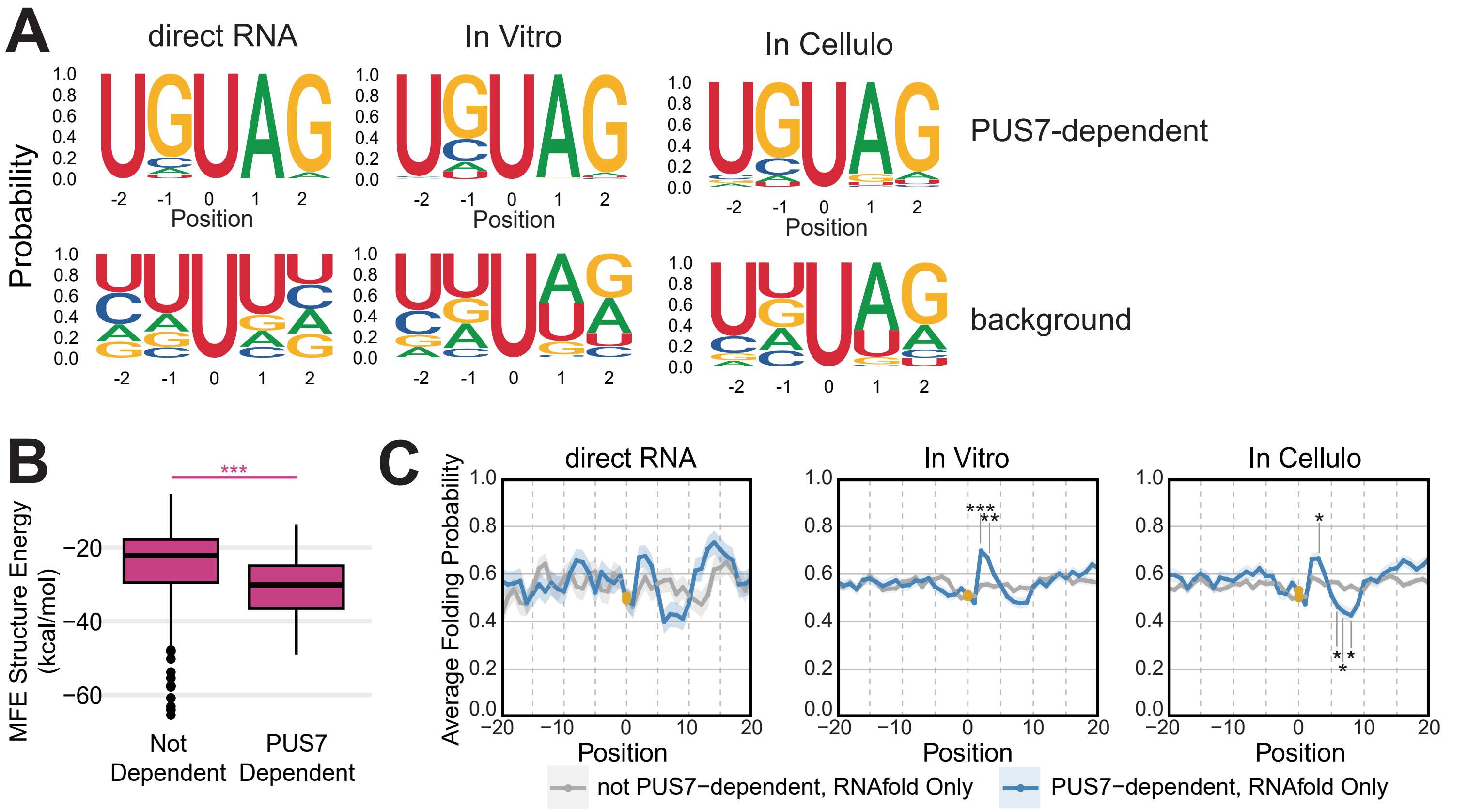


### Supplemental Figure 4

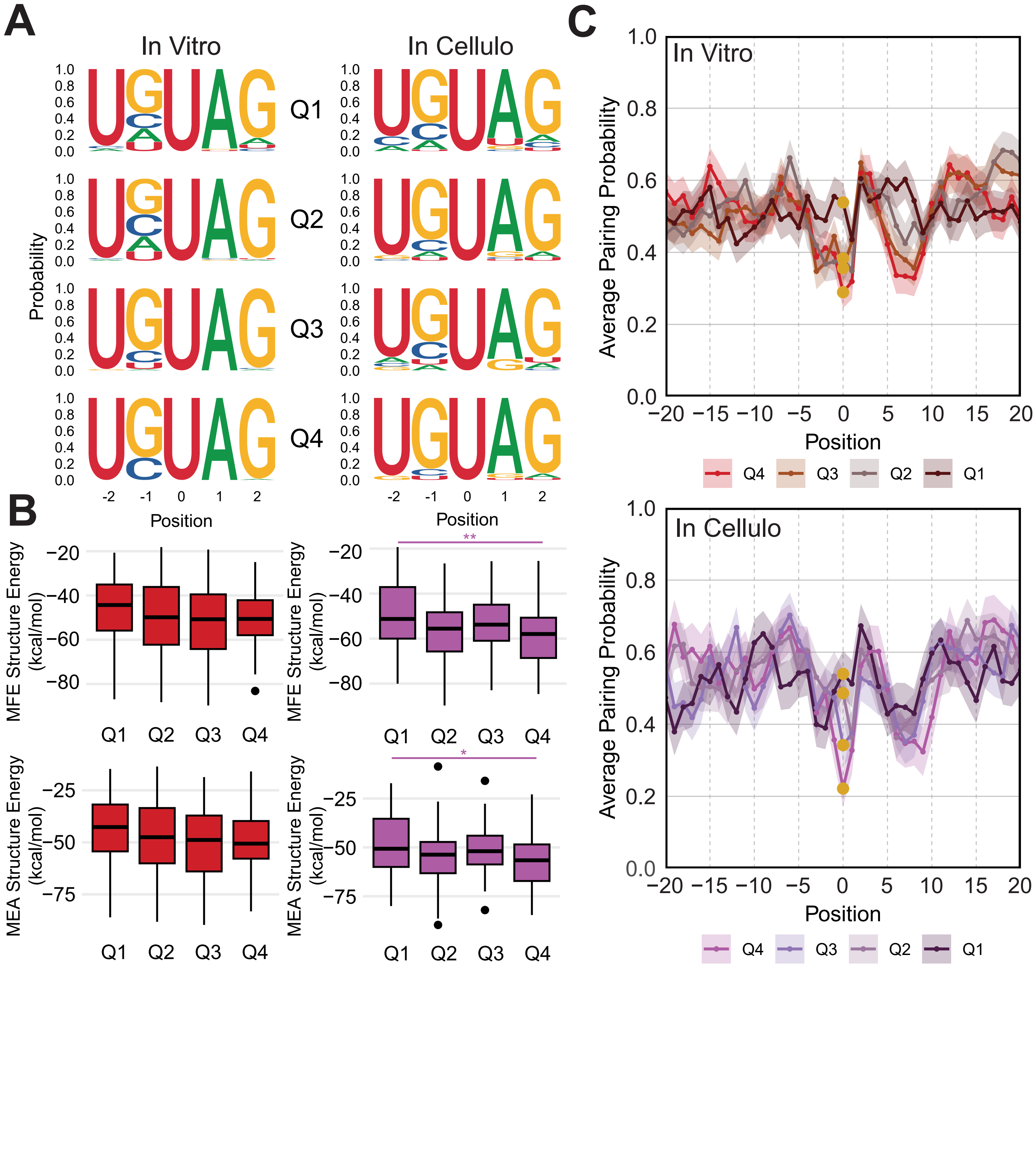


Supplemental Figure 5

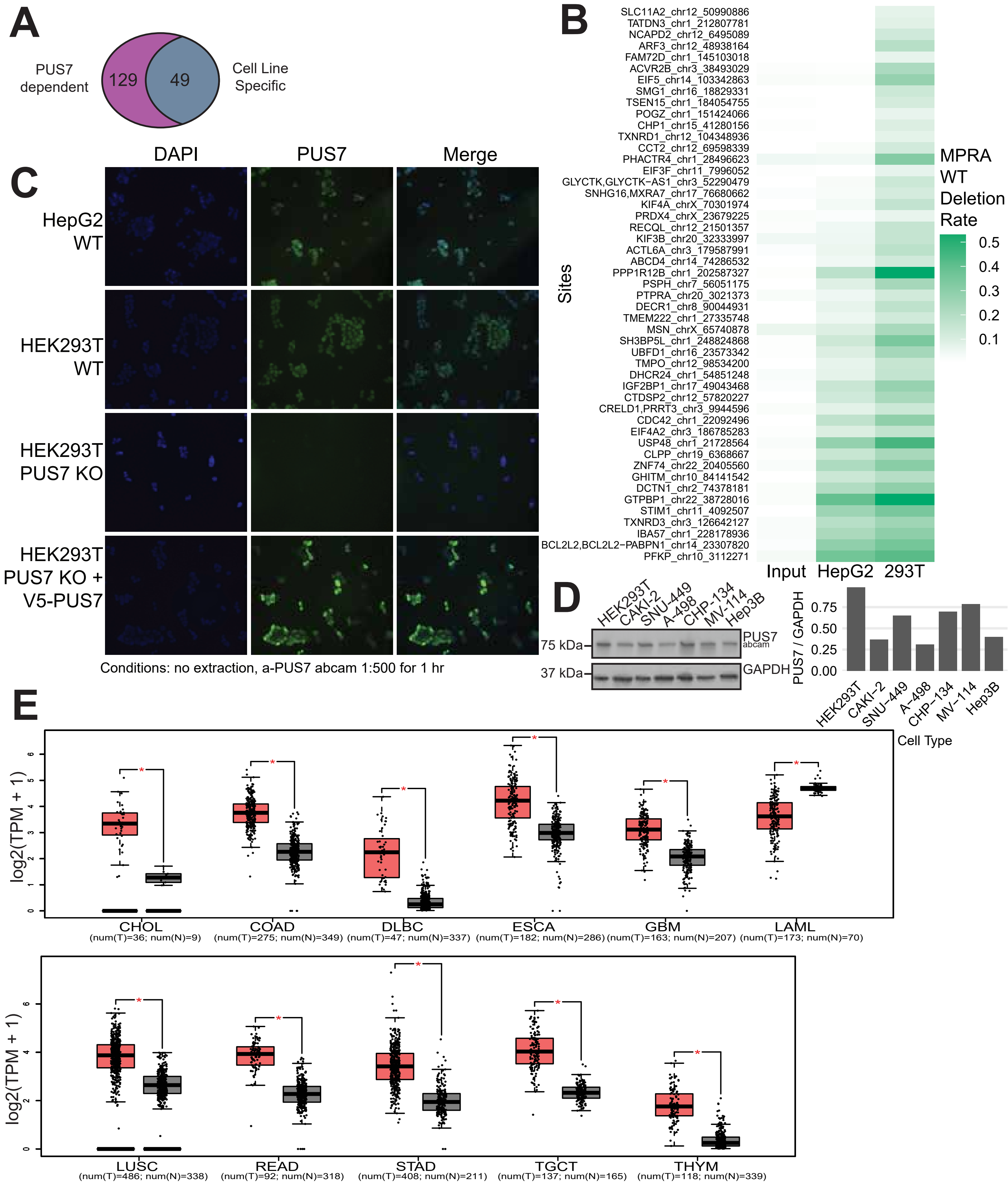


### Supplemental Figure 6
